# TET1 dioxygenase is required for FOXA2-associated chromatin remodeling in pancreatic beta-cell differentiation

**DOI:** 10.1101/2020.05.20.107532

**Authors:** Jianfang Li, Xinwei Wu, Minjung Lee, Jie Ke, Qingping Lan, Jia Li, Yun Huang, De-Qiang Sun, Ruiyu Xie

## Abstract

Existing knowledge of the role of epigenetic modifiers in pancreas development has exponentially increased. However, the function of TET dioxygenases in pancreatic endocrine specification remains obscure. We set out to tackle this issue using a human embryonic stem cell (hESC) differentiation system, in which *TET1*/*TET2*/*TET3* triple knock-out cells display severe defects in pancreatic β-cell specification. Integrative whole-genome analysis identifies unique cell-type-specific hypermethylated regions (hyper-DMRs) displaying reduced chromatin activity and remarkable enrichment of FOXA2, a pioneer transcription factor essential for pancreatic endoderm specification. Intriguingly, TET depletion leads to significant changes in FOXA2 binding at pancreatic progenitor stage, in which gene loci with decreased FOXA2 binding features low levels of active chromatin modifications and enriches for bHLH motifs. Transduction of full-length *TET1* but not the TET1-catalytic-domain in *TET*-deficient cells effectively rescues β-cell differentiation accompanied by restoring *PAX4* hypomethylation. Taking these findings together with the defective generation of functional β-cells upon TET1-inactivation, our study unveils an essential role of TET1-dependent demethylation in establishing β-cell identity. Moreover, we discover a physical interaction between TET1 and FOXA2 in endodermal lineage intermediates, which provides a new mechanistic clue regarding the complex crosstalk between TET dioxygenases and pioneer transcription factors in epigenetic regulation during pancreas specification.

## Introduction

During embryonic development, pluripotent human embryonic stem cells (hESCs) differentiate into many diverse lineages that make up the complex tissues and organs of the human body. Pancreatic lineage specification relies on crosstalk between the genome and environmental cues in the progenitor niche. This crosstalk is mediated by cis-regulatory elements that play a prominent role in spatiotemporal gene regulation during embryogenesis. In particular, distal regulatory elements, such as enhancers, serve as information integration hubs that allow binding of multiple regulators, including lineage-specific transcription factors (TFs) as well as epigenetic readers, writers, and erasers to ensure integration of intrinsic and extrinsic environmental cues at these loci ^1, 2^. It was recently demonstrated that pioneer TFs, such as FOXA1 and FOXA2, are required for proper chromatin opening and establishment of enhancer marks H3K4me1 and H3K27ac during pancreatic fate specification ^3, 4^. These pioneer TFs bind to nucleosomal DNA to initiate chromatin remodeling associated with DNA demethylation at newly accessible enhancers ^5–8^.

DNA demethylation is mediated by ten-eleven-translocation methylcytosine dioxygenases (TETs), which catalyze sequential oxidation of 5-methylcytosine (5mC) to 5- hydroxymethylcytosine (5hmC), 5-formylcytosine, and 5-carboxylcytosine ^9–13^. Distribution of 5hmC, a novel epigenetic modification, is dynamically changed by and positively correlated with active gene transcription during early lineage specification ^14–16^. Consequently, inhibition of TET family enzymes (TET1, TET2, and TET3) impairs cell fate commitment into neural, hematopoietic, cardiac, and several other lineages ^17–19^. We previously demonstrated that 5hmC positively correlates with ‘open’ chromatin at poised and active enhancers in multiple endodermal lineage intermediates using a stepwise hESC differentiation system toward pancreatic progenitors ^20^. However, the functional relevance and mechanisms by which TETs regulate pancreatic endocrine specification are currently unclear. While it is recently recognized that pioneer TF FOXA2 plays a critical role in enhancer priming during hESC pancreatic differentiation ^3, 4, 21^, there is still scarce understanding of whether and how FOXA2 interacts with TETs to mediate opening of surrounding chromatin.

## Results

### TET deficiency impairs pancreatic endoderm differentiation

To determine the biological significance of TET-dependent regulation during pancreas specification, we generated *TET1*, *TET2*, and *TET3* single knock-out (KO), double knock-out (DKO), and triple knock-out (TKO) H1 hESC lines using CRISPR/Cas9 technology (Supplementary Fig. 1a). Mutations resulted in premature termination codons, which were confirmed by Sanger sequencing. Global DNA hydroxymethylation levels from positive clones were assayed by 5hmC dot blot. A significant reduction of 5hmC signals was shown in TKO, TET1KO, TET1/2DKO, and TET1/3DKO cells, whereas TET2KO, TET3KO, and TET2/3DKO cells showed minimal alterations of 5hmC levels (Supplementary Fig. 1b). To avoid functional redundancy, we focused our initial analysis on TKO cells devoid of any TET-mediated active demethylation. Consistent with other *TET* knock-out mESC and hESC lines ^17, 22^, TKO H1-hESCs exhibited no apparent defects in stem cell self-renewal capacity or expression of pluripotent factors (Supplementary Fig. 1e).

To determine whether TET proteins affect hESC differentiation toward pancreatic endocrine fate, we used an established stepwise differentiation platform ^20^ to induce efficient differentiation of hESCs to definitive endoderm (DE), primitive gut tube (GT), pancreatic progenitor (PP), and pancreatic endocrine (PE) (Fig. 1a). Both TET triple-deficient lines (clones 2 and 6) displayed similar differentiation efficiency toward endoderm germ layer as the wild-type (WT) hESC line, in which over 90% of cells were SOX17^+^ by day 3 of differentiation (Supplementary Fig. 1c). Moreover, expression levels of other endoderm markers, such as *FOXA2*, *FOXA1*, and *CXCR4*, were unchanged in TET-depleted cells compared with control cells (Supplementary Fig. 1d, e). This analysis suggests that TET dioxygenases are dispensable for endoderm specification in the context of *in vitro* hESC differentiation. To examine the effects of TET ablation on pancreatic commitment, we subsequently examined the expression of critical pancreatic markers at the PP and PE stages. Using flow cytometry to quantitate the expression of *PDX1* at the PP stage and co-expression of *PDX1* and *NKX6.1* at the PE stage, we observed a substantial decrease in TET knock-out cells (Fig. 1b). These data were confirmed by immunofluorescence staining and RT-qPCR analysis (Fig. 1c, Supplementary Fig. 1e). Consistent with these findings, expression of the pancreatic progenitor markers *SOX9* and *PTF1A* and endocrine hormones insulin, and glucagon were significantly downregulated in TKO lines (Fig. 1c, Supplementary Fig. 1e). Interestingly, *PAX4*, a key determinant for β-cell specification, failed to induce in TET-inactivated cells, whereas expression of the α-cell determinant, *ARX*, was not inhibited (Supplementary Fig. 1e). These results demonstrate that TETs are required for pancreatic β-cell lineage specification.

**Fig. 1.**
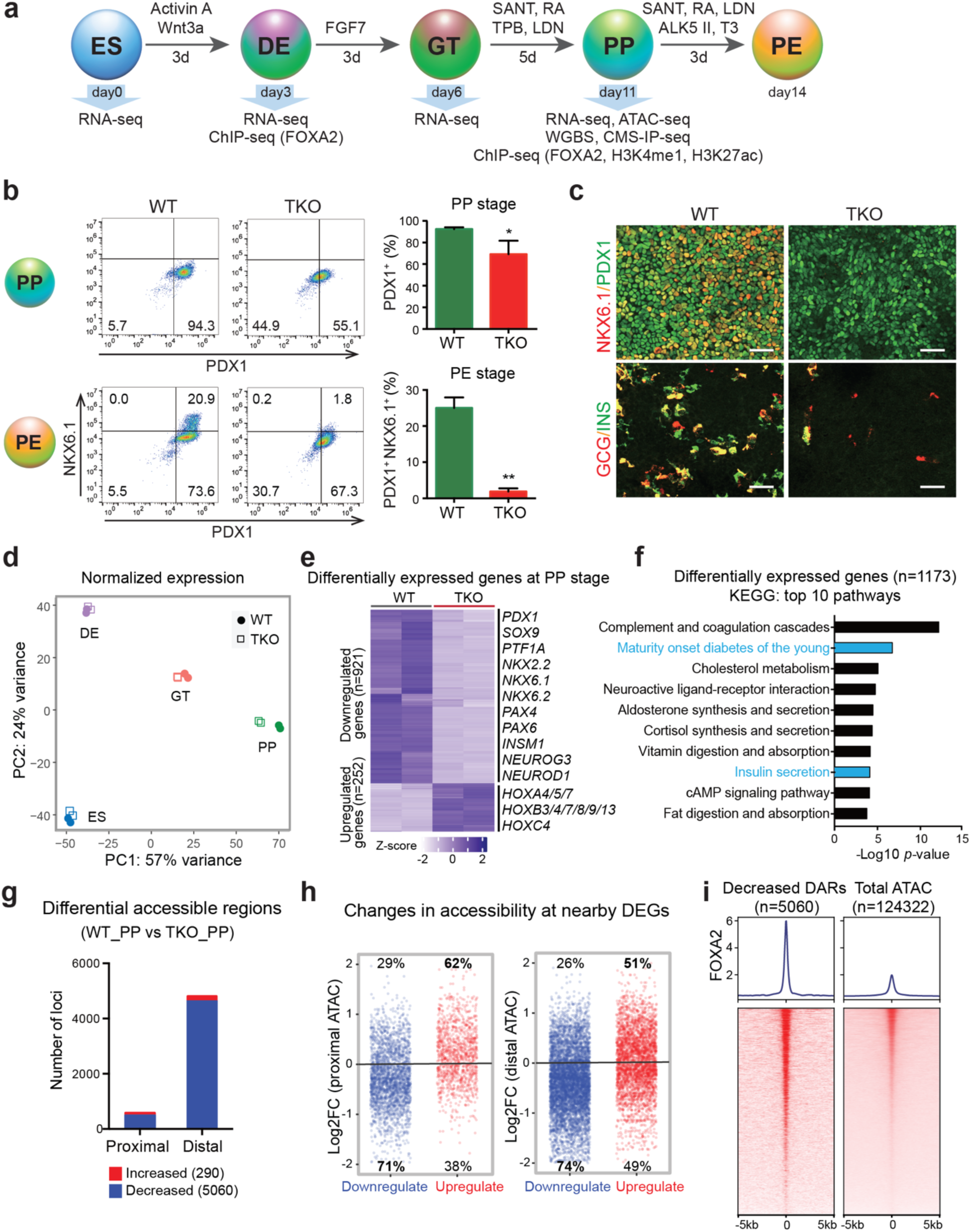
TET hydroxylases are required for pancreatic endocrine lineage specification. **a** Stepwise differentiation of hESCs to pancreatic endocrine cells. Wild-type (WT) and *TET1*/*TET2*/*TET3* triple knock-out (TKO) hESC lines were differentiated and analyzed by RNA-seq, WGBS, CMS-IP-seq, ATAC-seq, and ChIP-seq at the indicated stages (ES: embryonic stem cell; DE: definitive endoderm; GT: primitive gut tube; PP: pancreatic progenitor; PE: pancreatic endocrine). **b** Representative flow cytometry plots for PDX1 and NKX6.1 in WT or TKO cells at the PP and PE stages. Quantification of the percentage of PDX1^+^ NKX6.1^+^ cells is shown in the right panel. n = 3 independent differentiation. **c** Immunostaining of PDX1/NKX6.1 at the PP stage and insulin (INS)/glucagon (GCG) at the PE stage are shown in WT and TKO cells. Scale bar = 50 μm. **d** Principal component analysis showing variance in normalized transcriptome between WT and TKO cells at the ES, DE, GT, and PP stage. Each plotted point represents one biological replicate. **e** Genes with significant changes in expression in TKO cells relative to WT cells at the PP stage. Differentially expressed genes (DEGs) are classified into down- and upregulated groups. Each row corresponds to one individual gene and each column to a different biological replicate. The color scale from white to blue represents Z-score normalized gene expression levels from low to high (|fold change| ≥ 2; FDR < 0.05). **f** Functional analysis of DEGs in TKO_PP cells showing the top 10 KEGG pathways. Benjamini-Hochberg corrected *p*-values were used. **g** Number of regions with significant changes in chromatin accessibility upon TET depletion at proximal (≤ 1 kb from TSS) and distal (> 1 kb from TSS) regions (FDR < 0.05). **h** Dot plots depicting the ratios (WT_PP over TKO_PP) of ATAC-seq signals (y-axis) at proximal (left) or distal (right) regions of DEGs that were downregulated (blue) or upregulated (red) in TKO_PP cells compared with WT_PP cells. Values indicate the fraction of increased or decreased chromatin accessibility regions in down- and upregulated genes, respectively. **i** Average density plots (top) and heatmaps (bottom) of FOXA2 binding at decreased accessible regions (DARs) and total identified ATAC-seq peaks (± 5 kb). Heatmaps are ranked by decreased FOXA2 binding. The color scale from white to red represents the normalized signal from low to high.

To investigate TET-dependent global transcriptional changes during pancreatic endocrine cell fate commitment, we performed transcriptome analysis of WT and TKO cells at the ES, DE, GT, and PP stages. Principle component analysis of transcriptome data demonstrated that TET-deficient cells were similar at the ES, DE, and GT stages but differed substantially at the PP stage (Fig. 1d). Hierarchical clustering of genes specifically expressed at each stage revealed a remarkable downregulation of PP-specific genes in TKO_PP cells, whereas ES-specific and DE-specific genes showed no obvious changes in TET-deficient cells compared with WT cells (Supplementary Fig. 1f). In particular, 1,173 differentially expressed genes (DEGs) were identified in TKO_PP cells (Fig. 1e, Supplementary Table 1). Among them, there were 921 downregulated DEGs in TET-deficient cells, including many important developmental determinants for pancreatic endocrine specification, such as *PDX1*, *NKX2.2*, *NKX6.1*, *NKX6.2*, *NEUROG3*, and *PAX4*. We also observed 252 genes, particularly HOX family members, which were significantly upregulated upon TET deletion, suggesting that failure to differentiate into pancreatic β-cells is tightly linked to an aberrant TF network. In addition, KEGG pathway analysis of the DEGs showed enrichment for terms associated with maturity-onset diabetes of the young (MODY) and insulin secretion (Fig. 1f), implicating that loss of TETs impairs pancreatic endocrine formation at later stages of endoderm specification.

### FOXA2 binding is enriched at distal regulatory elements displaying decreased accessibility upon TET deletion

Given that our previous study demonstrated that TET-mediated cytosine oxidation is strongly associated with open chromatin regions during pancreatic differentiation ^20^, we used the assay of transposase-accessible chromatin followed by high-throughput sequencing (ATAC-seq) to evaluate chromatin accessibility landscapes in WT and TET-deficient cells. We assessed differential accessible regions (DARs) between WT_PP and TKO_PP cells from a total of 124,322 identified ATAC-seq peaks and found that 4.3% of ATAC-seq peaks significantly changed upon TET deletion (Supplementary Table 2). We found that loss of TET led to an overall reduction in chromatin accessibility, with 5,060 DARs showing reduced accessibility and only 290 DARs showing increased accessibility in TKO_PP cells (Supplementary Fig. 1g). Moreover, DARs with decreased accessibility were located primarily at distal regions (> 1 kb transcription start site (TSS)) (Fig. 1g). Decomposition of chromatin accessibility and DEGs in TKO_PP cells revealed that downregulated genes became less accessible (Fig. 1h).

To enhance the biological insights of TET depletion-mediated changes in chromatin accessibility, we analyzed the presence of TF binding motifs within DARs and found that regions with reduced accessibility were mostly enriched for motif of pioneer TF FOXA2 essential for pancreas organogenesis and chromatin remodeling ^4, 21, 23, 24^ (Supplementary Fig. 1h). Since the presence of a binding motif is not necessarily reflected of actual binding, we mapped FOXA2-binding sites by chromatin immunoprecipitation followed by next-generation sequencing (ChIP-seq) in pancreatic progenitors. Consistent with the motif prediction our analysis revealed that FOXA2 peaks coincide with ATAC-seq peaks preferentially decreased upon TET depletion (Fig. 1i). Collectively, our data indicate that failure to differentiate to pancreatic β-cells is likely to be linked with the loss of chromatin accessibility at distal regulatory elements enriched for pioneer TF FOXA2.

### Differentiation-specific DMRs feature reduced chromatin activity at pioneer TF binding sites

To investigate TET deficiency-mediated alterations in global DNA methylation, we performed whole-genome bisulfite sequencing (WGBS) in WT_PP and TKO_PP cells. Over 86% of sequencing reads were uniquely aligned to hg38 with a high sequencing depth of 24 *×* on CpG dinucleotides and typical bimodal distribution of methylation ratio in each sample (Supplementary Fig. 2a, b). The methylation ratio (5mC/C) was depleted at TSS but enriched across gene coding regions, especially within TKO_PP cells (Supplementary Fig. 2c). Among 26.6 million CpG sites detected in both WT_PP and TKO_PP cells (depth ≥ 5), a total of 251,658 differentially methylated cytosines (DMCs; credible difference of methylation ratio > 20%) were identified (Fig. 2a). Strikingly, 97.5% of DMCs were hypermethylated (hyper-DMCs), suggesting that TET inhibition results in a pronounced gain of methylation during pancreas differentiation. We observed enrichment of DMCs primarily at intergenic regions and introns (Supplementary Fig. 2d), whereas substantial enrichment of hypo-DMCs was also found in repeat elements, such as long interspersed elements, consistent with a suggested connection between hypomethylation and activation of transposable elements ^25, 26^. Based on an established link between DNA methylation and transcription regulation, we further calculated changes in methylation levels on cis-regulatory elements previously identified in pancreatic progenitors ^3, 27^. We found increased methylation at bivalent promoters, active enhancers, and, to a lesser extent, poised enhancers (Fig. 2b). Notably, hypermethylation was preferentially enriched at open chromatin (ATAC-seq) as well as FOXA2, GATA4 (GSE117136) ^4^, GATA6 (GSE117136) ^4^, and PDX1 (GSE117136) ^4^ distal binding regions (Fig. 2b, Supplementary Fig. 2e). This analysis demonstrates that active regulatory elements are hypermethylated upon TET depletion during pancreatic differentiation.

**Fig. 2.**
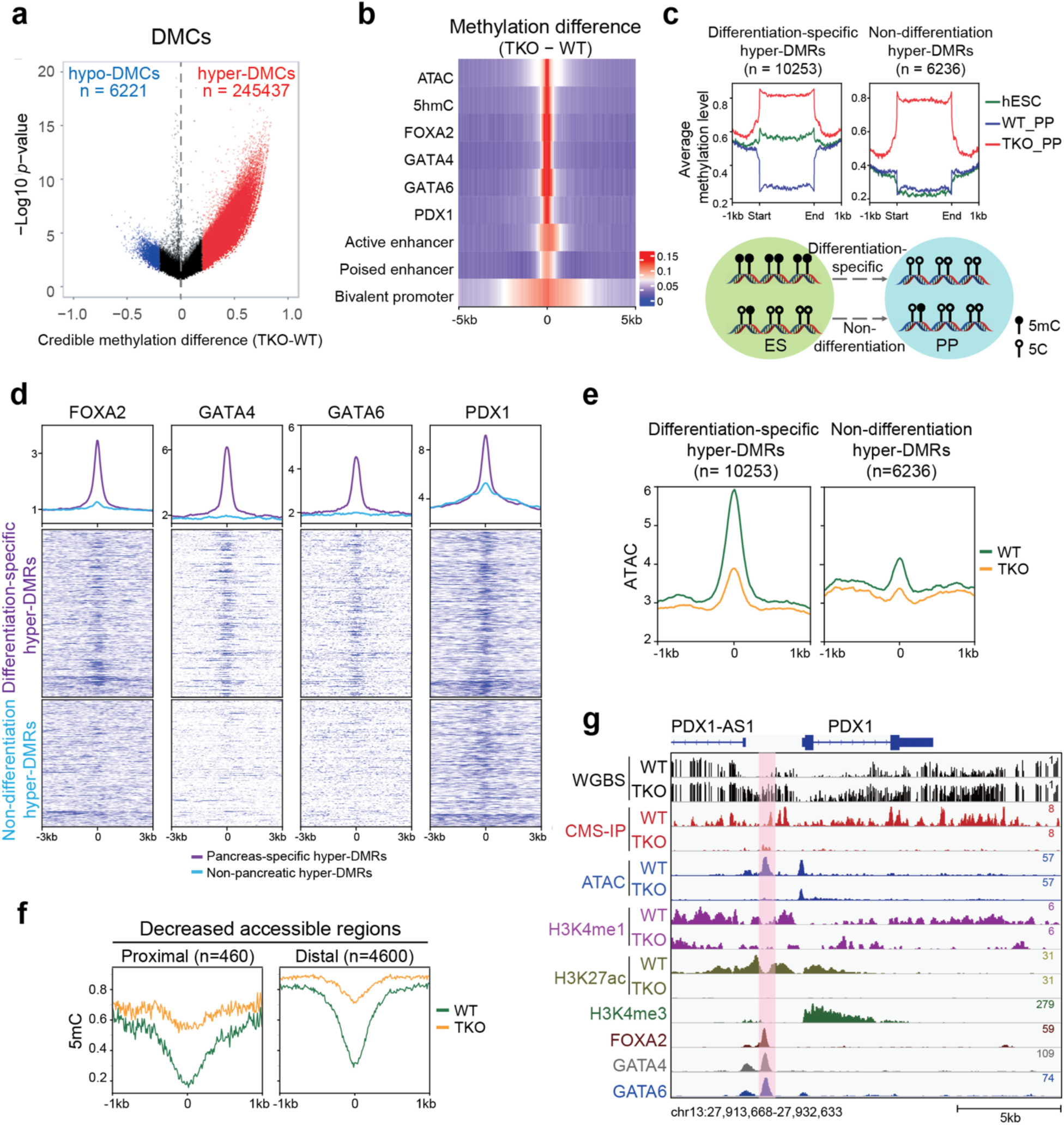
Differentiation-specific hyper-DMRs show reduced chromatin activity at pioneer TF binding sites. **a** Volcano plot of WGBS data illustrating differentially methylated CpGs (DMCs) identified in TKO_PP cells compared with WT_PP cells. Red and blue represent increased and decreased 5mC in TKO_PP cells, respectively (credible methylation difference > 0.2). **b** Heatmap illustrating methylation difference between TKO_PP and WT_PP cells at centers of annotated genomic features (± 5 kb) for chromatin accessibility (ATAC), hydroxylation (5hmC), TF binding (FOXA2, GATA4, GATA6, and PDX1), bivalent promoters, poised enhancers, and active enhancers. Average 5mC signals of every 100-bp bin were calculated. **c** Classification of TKO hyper-DMRs based on 5mC levels in hESCs (green), WT_PP cells (blue), and TKO_PP cells (red). **d** Average density plots and heatmaps of FOXA2, GATA4, GATA6, and PDX1 signals at differentiation-specific hyper-DMRs or non-differentiation hyper-DMRs in pancreatic progenitors. **e** Average density plots of ATAC-seq at differentiation-specific hyper-DMRs or non-differentiation hyper-DMRs in WT_PP (green) or TKO_PP (orange) cells. **f** Enrichment profile of methylation ratio (5mC/C) at proximal (≤ 1 kb from TSS) and distal (> 1 kb from TSS) decreased accessible regions in WT_PP (green) and TKO_PP (orange) cells. **g** Genome-browser view of the *PDX1*/*PDX1-AS1* locus. A specific TKO hyper-DMR showing decreased 5hmC, ATAC-seq, and H3K27ac signals is highlighted in pink.

Our previous studies reveal that DNA demethylation is correlated with pancreatic endocrine patterning ^20^. To gain better insight into the role of TET-mediated methylation changes in pancreatic differentiation, we analyzed differentially methylated regions (DMRs) by connecting at least three consecutive DMCs ^28^ between TKO_PP and WT_PP cells. We found a total of 16,490 hyper-DMRs and classified them into two categories based on their 5mC levels in pancreatic progenitor versus hESC ^29^ (Fig. 2c). Notably, more than half (n = 10,254) of hyper-DMRs exhibited decreased 5mC in WT_PP cells than in hESCs indicating an active demethylation process during pancreatic differentiation (named differentiation-specific hyper-DMRs), whereas the others (n = 6,236) showed similar methylation levels (non-differentiation hyper-DMRs) (Fig. 2c, Supplementary Fig. 2f). Interestingly, we found that pioneer TFs FOXA2, GATA4, GATA6, and PDX1 were predominantly enriched at differentiation-specific hyper-DMRs (Fig. 2d), suggesting that TET-dependent demethylation during pancreatic cell fate commitment occurs at loci primarily associated with lineage-specific TFs. We further compared ATAC-seq peaks at differentiation-specific hyper-DMRs with those at non-differentiation hyper-DMR. We found that chromatin accessibility was markedly reduced in TET-deficient cells at distally located hyper-DMRs in the differentiation-specific group (Fig. 2e, Supplementary Fig. 2g). Consistently, genomic loci displaying decreased accessibility showed a substantial increase in DNA methylation upon depletion of TET (Fig. 2f). For instance, the hyper-DMRs, which were annotated to *PDX1* and its lncRNA *PDX1-AS1* loci, overlapped with pioneer TF-binding sites and showed decreased chromatin activity in a TET-dependent manner (Fig. 2g). Significant hypermethylation was also found at the distal regulatory elements of *NEUROD1* and *NKX2.2* loci (Supplementary Fig. 2h). In summary, these analyses demonstrate that TET-dependent demethylation at TF-enriched distal regulatory elements is essential for chromatin remodeling during pancreas development.

### TET deficiency induces a differentiation-specific loss of 5hmC at FOXA2 target sites

To get additional insight into the TET-mediated hydroxymethylation network governing pancreatic differentiation, we used cytosine-5-methanesulfonate immunoprecipitation (CMS-IP) coupled with high-throughput sequencing to profile 5hmC landscapes in WT_PP and TKO_PP cells. As a consequence of TET deletion, we found genome-wide loss of hydroxymethylation in TET-deficient cells (Supplementary Fig. 3a, b). As expected, 94% of CMS-positive regions showed a significant reduction in TKO_PP cells relative to WT_PP cells (Supplementary Fig. 3c). These differentially hydroxymethylated regions displaying reduced 5hmC (hypo-DHMRs) were primarily located at non-promoter regions (> 1 kb from TSS) (Supplementary Fig. 3d).

Reduced DNA hydroxymethylation in TET-deficient cells could be a result of the initial loss of 5hmC at the ES stage or unsuccessful oxidation of 5mC during lineage progression. To provide biological insight into differentiation-specific changes of hydroxymethylation upon TET inactivation, we systematically analyzed hypo-DHMRs overlapped with previously identified regions showing de-novo increased 5hmC during hESC pancreatic differentiation ^20^. Specifically, 27,113 hypo-DHMRs with higher 5hmC levels in WT_PP than in WT_ES (named ‘differentiation-specific hypo-DHMRs’) were first isolated from a total of 70,857 TKO hypo-DHMRs (Fig. 3a). We subsequently clustered them into four categories based on the dynamic changes of 5hmC across stages in WT cells (Fig. 3b). In particular, one group gained 5hmC specifically at the PP stage (PP-specific), whereas others gained 5hmC at the earlier DE (DE-to-PP) or GT (GT-to-PP) stages and sustained hydroxymethylation until the PP stage. We found that GATA motifs were mostly associated with the DE-to-PP cluster, in which gain of 5hmC began at the DE stage (Fig. 3c, purple).

**Fig. 3.**
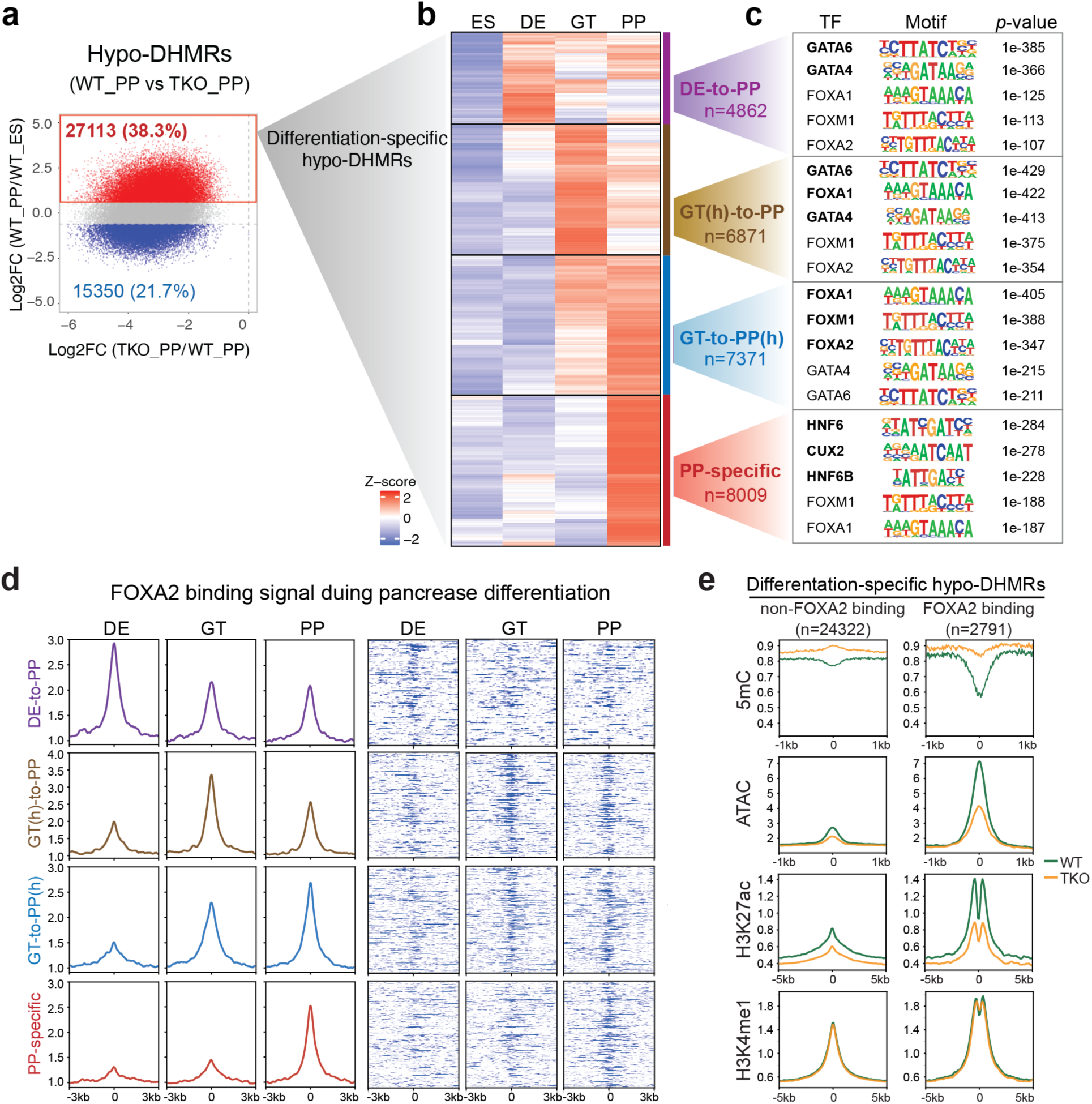
Loss of TET disrupts epigenetic dynamics at differentiation-specific hypo-DHMRs associated with distinct pioneer TFs motifs. **a** Scatterplot showing log_2_-fold change of 5hmC in pancreatic progenitors (WT_PP) versus hESCs (WT_ES) within hypo-DHMRs identified in TKO_PP cells. Red indicates increased 5hmC and blue shows decreased 5hmC in pancreatic progenitors relative to hESCs (|fold change| ≥ 1.5, FDR < 0.05). **b** Clustering of 5hmC signals in WT cells at ES, DE, GT, and PP stage. Each row represents one hypo-DHMR showing increased 5hmC in pancreatic progenitors compared with hESCs. Red represents high 5hmC; blue indicates low 5hmC. **c** Motif enrichment analysis among the four clusters defined in **b**. The top five known motifs are shown. **d** Average density plots and heatmaps showing dynamic changes of FOXA2 binding in DE, GT, and PP lineage intermediates across the four clusters defined in **b**. The color scale from white to blue in heatmaps represents normalized FOXA2 signal from low to high. **e** Enrichment profile of methylation ratio (5mC/C), chromatin accessibility (ATAC), H3K27ac, and H3K4me1 at ‘differentiation-specific hypo-DHMRs’ alone (left column) and ‘differentiation-specific hypo-DHMRs’ overlapped with FOXA2-bound sites (right column). The WT signal is marked in green and the TKO signal in orange.

By contrast, the PP-specific cluster displayed a prominent presence of binding sites for HNF6, which is critical for pancreatic endocrine differentiation ^30^ (Fig. 3c, red). Most notably, FOXA motifs were predominantly associated with the GT-to-PP(h) cluster in which 5hmC levels increased at the GT stage and continued to be elevated until the PP stage (Fig. 3c, blue), whereas both FOXA and GATA motifs were highly associated with the GT(h)-to-PP cluster in which 5hmC peaked at the GT stage and subsequently decreased at the PP stage (Fig. 3c, brown). Collectively, these results suggest a unique temporal binding pattern of GATAs, FOXAs, and HNF6s associated with the dynamic distribution of 5hmC during pancreatic differentiation, which is supported by a recent study demonstrating sequential requirements for these TFs during transitions in pancreas development ^4^.

### 5hmC and FOXA2 co-occupy target sites are essential for chromatin opening and enhancer activation

Given that hydroxymethylation is positively associated with chromatin activity ^20^ and FOXA2 is strongly enriched in regions featured decreased chromatin accessibility in TET-deficient cells (Fig. 1i), we wondered if binding of FOXA2 dynamically correlated with TET-mediated hydroxymethylation. Thus, we mapped FOXA2-binding sites by ChIP-seq at the DE, GT, and PP stages where FOXA2 continually expressed. Consistent with the motif predictions our analysis revealed that FOXA2 deposition corresponded well with the dynamic changes of 5hmC across differentiation stages in the four ‘differentiation-specific hypo-DHMRs’ clusters (Fig. 3d). Together, these data support that FOXA2 binding is highly dynamic during pancreatic differentiation and strongly prefers differentiation-associated 5hmC deposition sites.

It was previously demonstrated that the pioneer TF FOXA2 is essential for enhancer priming during pancreatic differentiation ^3, 4^. We, thus, speculated that loss of TETs inhibits enhancer activation particularly at hypo-hydroxymethylated regions enriched with FOXA2 binding. We conducted ChIP-seq for enhancer signatures H3K4me1 and H3K27ac in WT_PP and TKO_PP cells and performed integrated analysis. We found a remarkable decrease in ATAC-seq and H3K27ac signal at ‘differentiation-specific hypo-DHMRs’ overlapped with FOXA2 peaks, where DNA methylation was dramatically increased upon TET depletion (Fig. 3e). Nevertheless, FOXA2 and other pioneer TFs expressed at similar levels in TKO and WT cells (Supplementary Fig. 3e), implicating that inhibition of TET did not alter their expression. Collectively, these data reveal that TET deficiency results in a progressive failure of 5mC oxidation at a subset of FOXA2 targets essential for the establishment of active enhancers during differentiation.

### FOXA2 transiently binds to differentiation stage-specific genomic loci enriched for bHLH motifs

Being a pioneer TF, FOXA2 is expected to engage its binding sites on nucleosomal DNA to mediate nucleosome depletion and chromatin remodeling ^6, 31^. Despite its potentially universal binding ability, FOXA2 targeting was highly dynamic between pancreatic differentiation states (Fig. 3d). We, therefore, ask if additional factors can influence the binding of FOXA2. We tackled these questions by mapping FOXA2 binding sites in WT and TKO cells at the DE and PP stages, respectively. Consistent with previous reports ^4, 32^, FOXA2 was primarily located at non-promotor regions (Supplementary Fig. 4a) implicating that FOXA2 is mainly involved in transcription regulation through distal regulatory elements. Upon TET depletion, 99.5% FOXA2 binding sites did not show changes at the DE stage (Fig. 4a), while FOXA2 occupancy was dramatically changed at the PP stage, in which 10% and 6% of FOXA2 target sites showed reduced and greater FOXA2 binding, respectively (Supplementary Fig. 4b, Supplementary Table 3). Since DNA methylation and hydroxymethylation status were significantly altered in TET-deficient cells (Fig. 2a, Supplementary Fig. 3c), we wondered whether increases in methylation or decreases in hydroxymethylation contributed to differential recruitment of FOXA2 in TKO_PP cells. We first assessed changes in FOXA2 binding in WT_PP versus TKO_PP cells at the hyper-DMRs or hypo- DHMRs identified in TKO_PP cells. We found that ∼75% regions showed no changes in FOXA2 binding (Fig. 4b). We then tested if only hypermethylated or hypo-hydroxymethylated FOXA2- bound sites would be affected. We found that similar proportions of FOXA2 target sites displayed reduced/greater of FOXA2 binding regardless of the presence or absence of hypermethylation (Fig. 4c). Similarly, hypo-hydroxymethylated FOXA2-bound sites did not show a preference in gain/loss of FOXA2 binding compared to iso-hydroxymethylated FOXA2-bound sites (Fig. 4d). Consistent with the ability of pioneer TFs to engage with inaccessible chromatin, our results strongly demonstrate that DNA methylation/hydroxymethylation states do not interfere in FOXA2 binding.

**Fig. 4.**
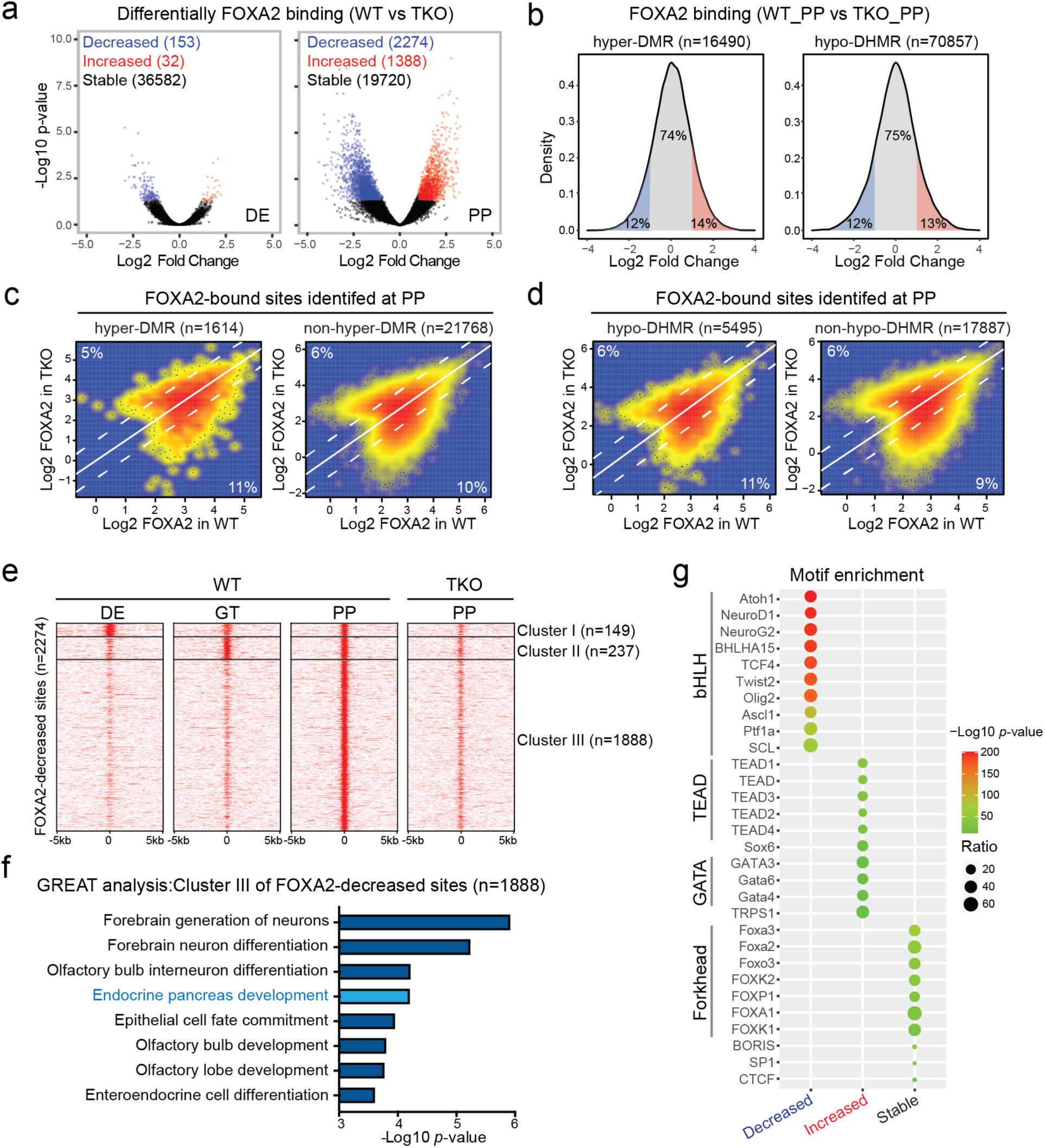
FOXA2 transiently binds to differentiation stage-specific genomic loci enriched for bHLH motifs. **a** Volcano plots of FOXA2 ChIP-seq data illustrating differential FOXA2 binding sties identified in TKO cells compared to WT cells at the DE (left) and PP stage (right). Red and blue represent increased and decreased FOXA2 signal in TKO cells, respectively (|fold change| ≥ 2; FDR < 0.05). **b** Density plots showing the percentages of differential FOXA2 binding in TKO_PP cells compared to WT cells at hyper-DMRs (left) and hypo-DHMRs (right). The color of red, blue, and gray represents increased, decreased, and non-changed FOXA2 signals in TKO_PP cells, respectively (|fold change| ≥ 2). **c** Scatterplots presenting FOXA2 signals in WT_PP cells versus TKO_PP cells at FOXA2 binding sites overlapped with (left) or without (right) hyper-DMRs. Percentages of differential FOXA2 binding are indicated (|fold change| ≥ 2). **d** Scatterplots presenting FOXA2 signals in WT_PP cells versus TKO_PP cells at FOXA2 binding sites overlapped with (left) or without (right) hypo-DHMRs. Percentages of differential FOXA2 binding are indicated (|fold change| ≥ 2). **e** Clustering of FOXA2 binding signals at the DE, GT, and PP stage in regions showed lost FOXA2 in TKO_PP cells. **f** Pathway enrichment annotations from GREAT for cluster III decreased FOXA2-bound sites. Benjamini-Hochberg corrected *p*- values were used. **g** Transcription factor motif enrichment analysis of genomic regions showed decreased, increased, and stable FOXA2 binding in TKO_PP cells. The top 10 known motifs are shown. The color representing *p*-values and the size of circle representing proportion of peaks containing a TF motif for each group.

To comprehensively characterize temporal patterns of FOXA2 recruitment at the FOXA2- decreased binding sites, we clustered FOXA2 binding signal in WT cells at the DE, GT, and PP stages. Three distinct patterns of FOXA2 occupancy were observed. Cluster I regions (149) were bound by FOXA2 from DE to PP stage, cluster II regions (237) were FOXA2-bound at GT and PP stages, while the most predominant group, cluster III (1888), displayed de novo FOXA2 occupancy in pancreatic progenitors (Fig. 4e). We then analyzed the annotations of nearby genes with Genomic Regions Enrichment of Annotations Tool (GREAT) and found that cluster III regions were enriched for terms of neuron and endocrine pancreas development (Fig. 4f). Further examination of ATAC-seq, H3K27ac, and 5mC signals in three FOXA2-decreased clusters revealed high levels of DNA methylation accompanied by low levels of enhancer activity and chromatin accessibility in cluster III regions (Supplementary Fig. 4c). Taken together, our data imply that a subset of FOXA2 transiently binds to differentiation stage-specific genomic loci with low levels of active chromatin modifications at which additional lineage-specific TFs may be required to facilitate FOXA2 binding and subsequent chromatin remodeling.

To identify potential TFs associated with differential FOXA2 binding we determined unique DNA binding motifs within FOXA2-decreased, increased and stable sites. Interestingly, loci lost FOXA2 binding, particularly in cluster III, were mostly enriched for basic-helix–loop–helix (bHLH) motifs such as the pancreatic endocrine cell fate determinant NEUROD1 ^33^ (Fig. 4g, Supplementary Fig. 4d). In contrast, regions gained FOXA2 were mostly enriched with motifs of TEAD and GATA family members, and FOXA2-stable sites most prominently feature forkhead and CTCF motifs (Fig. 4g). Notably, expression of the bHLH TFs including *NEUROD1*, *PTF1A*, and *ASCL1* failed to be induced in TET-knockout cells (Supplementary Fig. 4e) which likely results in a decrease of FOXA2 binding at genomic loci primarily associated with these TFs. In summary, these data suggest that recruitment of FOXA2 to differentiation stage-specific sites is genetically and epigenetically primed with the cooperation of additional lineage-specific TFs.

### TET1 is required in pancreatic β-cell specification

Our findings suggest that TET inactivation-induced aberrant methylation/hydroxymethylation at least in part contributes to defective *β*-cell specification. To determine the essential role of each TET family member in pancreatic differentiation, we further analyzed mutants with single (*TET1*, *TET2*, or *TET3*) or double (*TET1/2*, *TET2/3*, or *TET1/3*) *TET* knock-out. All mutant lines were able to induce the expression of *PDX1* and *NKX6.1* to levels comparable to WT (Supplementary Fig. 5a, b). However, only those retaining intact *TET1* expression (TET2KO, TET3KO, and TET2/3DKO) displayed proper induction of *PAX4* (Supplementary Fig. 5c). In comparison with *TET2*/*TET3* double-deletion, inhibition of TET1 alone had significant effects on the formation of INS- and C-peptide (CPEP)-expressing *β*-cells but not GCG-expressing *α*-cells (Fig. 5a, b).

**Fig. 5.**
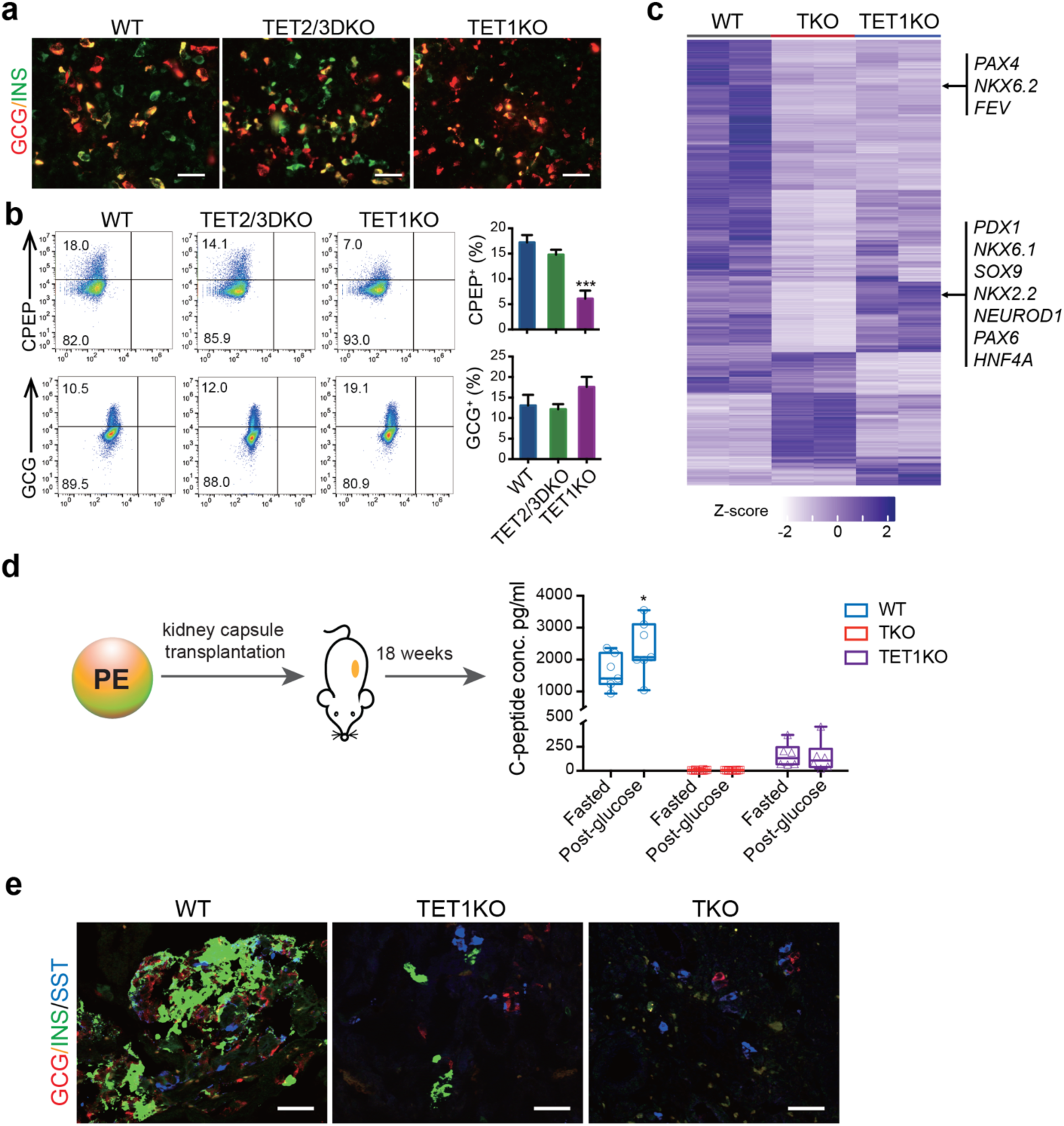
TET1 knock-out cells show impaired differentiation into functional β-cells. **a** Immunostaining of INS, and GCG in WT, TET2/3DKO, and TET1KO cells at the PE stage. Scale bar = 50 μm. **b** Representative plots of flow cytometry of human C-peptide (CPEP) and glucagon (GCG) in WT, TET2/3DKO, and TET1KO cells at the PE stage. Quantifications of the percentage of CPEP^+^ or GCG^+^ cells are shown in the right panel. n = 3 independent differentiation. **c** Heatmap showing the hierarchical clustering of DEGs among WT, TKO, and TET1KO cells at the PP stage. Each row represents one DEG, and each column represents one biological replicate. The color scale from white to blue represents normalized gene expression levels from low to high (|fold change| ≥ 2; FDR < 0.05). **d** WT, TKO, and TET1KO cells were differentiated to the PE stage and transplanted under the kidney capsule of nondiabetic female SCID-beige mice. Eighteen weeks post-implantation, human C-peptide levels were measured after an overnight fast and 30 min following an i.p. glucose injection. C-peptide levels from individual mice are shown in box and whisker plots. **e** Immunostaining of insulin (INS), glucagon (GCG), and somatostatin (SST) in cell grafts from WT_PE, TET1KO_PE, and TKO_PE transplanted mice. Scale bar = 50 µm.

To evaluate global transcriptome changes in response to *TET1* deletion, we performed RNA-seq of TET1KO cells at the PP stage. Integrated analysis of WT_PP, TKO_PP, and TET1KO_PP data revealed a total of 1,590 DEGs among the three lines (Supplementary Table 4). Specifically, 555 genes, including most pancreas developmental regulators such as *PDX1*, *SOX9*, *NKX2.2*, *NKX6.1*, *NEUROD1*, and *PAX6*, were downregulated in TKO_PP cells but not in TET1KO_PP cells (Fig. 5c), consistent with effective differentiation of TET1KO-hESCs into PDX1^+^ cells (Supplementary Fig. 5a). However, 536 genes, including the *β*-cell fate determinants *PAX4*, *NKX6.2*, and *FEV* ^34–36^, were significantly inhibited in TET1-deficient cells, implicating TET1 as responsible for the activation of a subset of genes essential for *β*-cell identity. To further analyze the functional consequences of TET1 inhibition, WT-, TET1KO-, and TKO-hESCs were differentiated to the PE stage and subsequently engrafted into SCID-beige mice under the kidney capsules (Fig. 5d). Glucose-stimulated human C-peptide secretion was determined 18 weeks post-transplantation. Notably, mice engrafted with WT_PE cells produced substantial fasting C-peptide in serum (1605 ± 527 pg/ml) and showed statistically significant glucose-stimulated C-peptide secretion (2,358 ± 839 pg/ml). By contrast, mice transplanted with TET1KO_PE cells secreted extremely low amounts of basal C-peptide (163 ± 122 pg/ml) and showed no response to glucose stimulation (149 ± 161 pg/ml). In agreement with these functional results, excised WT_PE cell grafts were highly composed of insulin^+^ *β*-cells (Fig. 5e). The TET1KO_PE cell grafts displayed much less insulin content, and only the *δ*-cell hormone somatostatin was detected in TKO_PE cell grafts. Collectively, these data demonstrate that loss of TET1 impairs *β*-cell specification and maturation.

### Full-length TET1 is required for the *PAX4* enhancer to achieve a hypomethylated state

To further determine whether TET1 is responsible for *β*-cell specification, we restored the full-length (*TET1FL*) as well as the *TET1*-catalytic-domain (*TET1CD*) expression, respectively, using lentivirus-mediated gene transduction in TKO cells. In contrast to the catalytically inactive *TET1* (*TET1CD^mut^*) (H1672Y, D1674A), both *TET1FL-* and *TET1CD*-transduced cells showed a global increase in 5hmC levels, confirming the replenishment of hydroxymethylation (Supplementary Fig. 6a). Differentiation of the TKO-TET1FL line toward pancreatic endocrine fate led to robust induction of *β*-cell determinants, including *PDX1*, *NKX6.1*, and *PAX4*, whereas transduction of *TET1CD* only resulted in a slight increase of *NKX6.1* and *PAX4* (Fig. 6a, b, Supplementary Fig. 6c). FACS analysis and immunostaining further demonstrated that TET1CD-transduced cells showed an effective generation of GCG-producing *α*-cells but not INS/CPEP-expressing *β*-cells at the PP stage (Supplementary Fig. 6b, c), indicating that full-length TET1 is critical for *β*-cell specification.

**Fig. 6.**
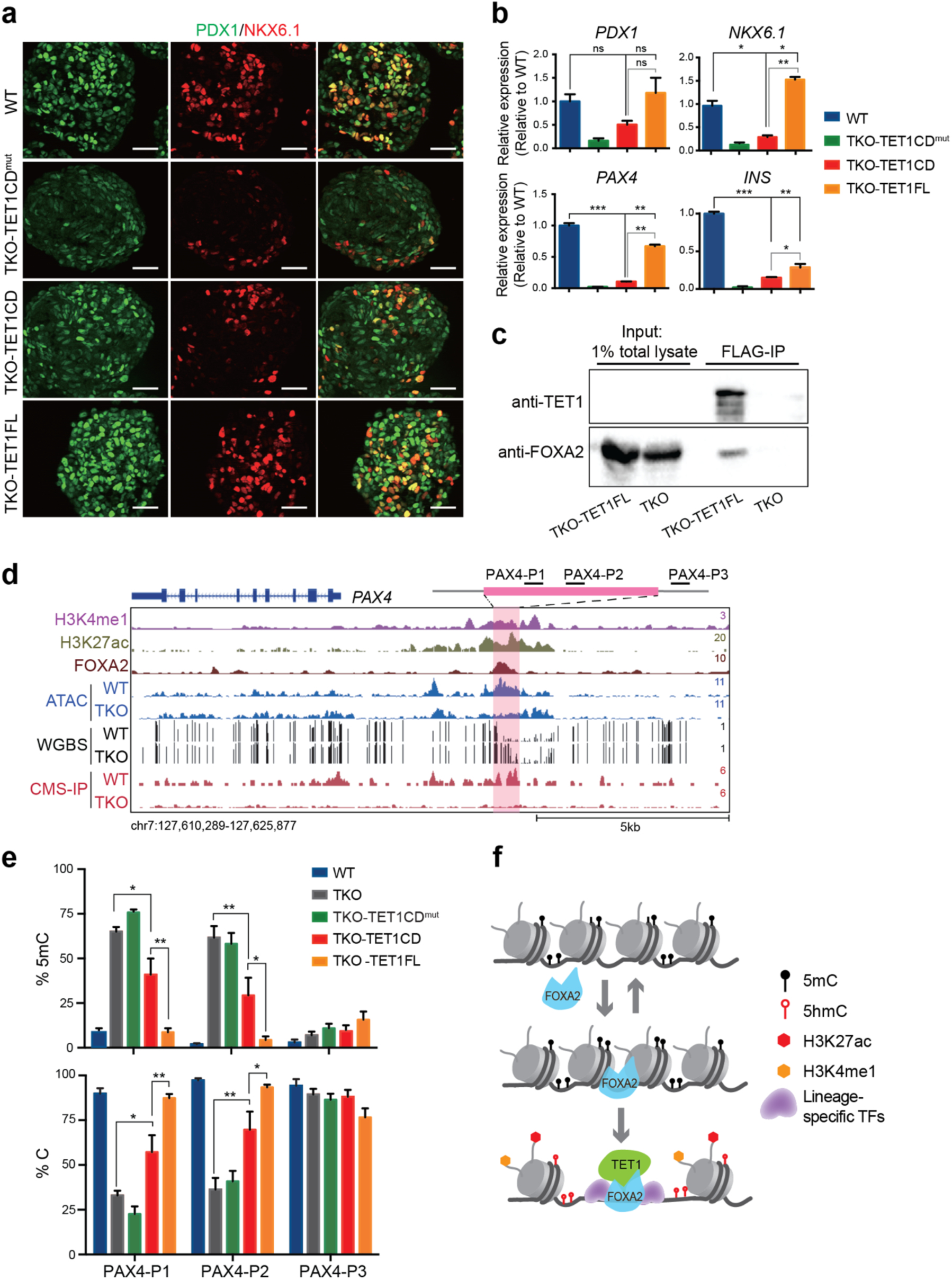
Full-length TET1 is required for the establishment of a hypomethylated PAX4 enhancer. **a** Immunostaining of PDX1 and NKX6.1 at the PP stage for WT, TKO-TET1CD^mut^, TKO-TET1CD, and TKO-TET1FL cells. Scale bar=50 μm. **b** Expression of *PDX1*, *NKX6.1*, *PAX4*, and *INS* in WT, WT, TKO-TET1CD^mut^, TKO-TET1CD, and TKO-TET1FL cells at the PE stage. **c** Western blots showing immunoprecipitation with antibodies against FLAG on cell extract of TKO-TET1FL cells followed by immunoblotting with antibodies against TET1 or FOXA2. TKO is included as control. **d** Genome-browser view of the *PAX4* locus with increased methylation and decreased chromatin associability upon TET depletion at a FOXA2-bound site featured enhancer signatures H3K4me1 and H3K27ac. **e** Locus-specific increase in 5mC at the *PAX4* enhancer in TKO or TKO-TET1CD samples compared with WT and TKO-TET1FL samples. Percentages of unmethylated cytosine and 5mC at CCGG sites are shown. n = 3 independent differentiation. **f** Schematic model depicting cooperative interaction among TET1 and TFs in lineage-specific enhancer activation. Pioneer TF FOXA2 transiently binds to enhancers. Lineage-specific TFs, such as PTF1A and NEUROD1, stabilize the binding of FOXA2 which recruits TET1 to induce DNA demethylation and subsequent chromatin opening.

As TET1 depletion inhibited the induction of genes essential for *β*-cell specification (Fig. 5), we reasoned that co-binding of TET1 and FOXA2 at distal regulatory elements modulates focal methylation status and subsequent gene activation. To identify potential interaction between TET1 and FOXA2, we performed immunoprecipitation in TKO-TET1FL cells expressing a FLAG-tagged *TET1FL* gene. Anti-FLAG antibodies co-precipitated endogenous FOXA2 in TKO-TET1FL cells differentiated at the DE stage (Fig. 6c). In contrast, co-precipitation of other endoderm proteins, including SOX17, GATA6, and FOXA1, was not observed (Supplementary Fig. 6d). These data illustrate that TET1 physically interacts with FOXA2 and therefore, can be recruited to FOXA2-bound loci to mediated demethylation during lineage induction.

Since the induction of *PAX4* was dramatically inhibited in TET1-deficient lines (Supplementary Fig. 5b), we asked if TET1 would cooperate with FOXA2 to mediate focal demethylation at the *PAX4* enhancer. We found that a putative *PAX4* enhancer (∼4.0 kb upstream of the TSS), where FOXA2 bound and displayed hypermethylation in TET-deficient cells (Fig. 6d, pink area). We next examined whether the elimination of TET1 alone results in hypermethylation at this site. Locus-specific methylation was determined using a glucosylation and digestion-based method followed by qPCR analysis. Two independent pairs of primers (PAX4-P1 and PAX4-P2) located within the FOXA2-binding site (Fig. 6d) were used to determine the percentage of methylated and non-methylated cytosine in TET1KO, TET2/3DKO, TKO, and WT cells at the PP stage. Notably, the percentage of 5mC was increased to 35–50% for TET1KO and 60–65% for TKO compared with WT but remained unchanged for TET2/3DKO (Supplementary Fig. 6e, left panel). A corresponding decrease in unmethylated cytosine content was observed, with values of 45–65% and ∼35% of C in TET1KO and TKO samples, respectively (Supplementary Fig. 6e, right panel). As a housekeeping control, we examined methylation content at a nearby region using the PAX4-P3 primer, and as expected, no significant differences in 5mC and C were observed in any samples. We then determined methylation contents at the same *PAX4* locus in TET1-replenished TKO cells. Consistent with the results found in TET2/3DKO cells, no detectable differences in 5mC and C were found between TKO-TET1FL and WT samples (Fig. 6e). However, 5mC levels were significantly higher with a value of 30-40% in TKO-TET1CD than TKO-TET1FL samples (Fig 6e, top panel). Together with the insufficient restoration of *PAX4* in TKO-TET1CD cells (Fig. 6b), we conclude that full-length TET1 is responsible for the establishment or maintenance of a hypomethylated state at the *PAX4* enhancer, which is essential for transcription activation of *PAX4* and subsequent *β*-cells generation.

## Discussion

In the present work, we dissected the roles of TET proteins in pancreatic endocrine commitment based on a stepwise hESC differentiation system. We found that the loss of all three TET family members significantly impaired the differentiation of pancreatic *β*-cells. Furthermore, we discovered that locus-specific hypermethylation was associated with genes essential for *β*-cell specification and maturation, such as *PAX4* ^34^, *PDX1* ^37^, and *NKX2.2* ^38^. The reintroduction of *TET1* in TET-deficient cells effectively reversed hypermethylation and restored the expression of *PAX4*. We further demonstrated that TET1 functions as an upstream epigenetic regulator of *PAX4* through direct recruitment by FOXA2 to a putative *PAX4* enhancer to preserve its unmethylated status, thereby potentiating *PAX4* expression to adopt β-cell fate during endocrine lineage commitment. Consistently, we observed striking increases in methylation at the *PAX4* enhancer in TET1 knock-out cells but not in TET2/TET3 DKO cells, suggesting that TET1 epigenetically regulates induction of the *β*-cell program in a locus-specific manner. Moreover, despite the successful induction of *PDX1* and *INS* upon deletion of TET1 alone, mice receiving TET1KO cell grafts showed a persistent defective insulin response to glucose, implying that TET1 is also essential for *β*-cell maturation.

Whereas the specification of *β*-cells was strongly influenced by depletion of *TET1*, we did not observe a significant inhibition on the expression of *ARX*, a critical *α*-cell fate determinant ^39^. In contrast to *PAX4*, no hyper-DMRs were identified within the *ARX* locus, where methylation levels were nearly undetectable in pancreatic progenitors. Previous studies demonstrate that several CpG-rich sites of *Arx*, including one site close to TSS and another site 2 kb upstream of TSS, are heavily methylated in adult *β*-cells but not *α*-cells ^40^. Moreover, pharmacological inhibition of DNA methyltransferases, *Dnmts*, in pancreatic progenitors promotes α-cell specification ^41^, whereas deletion of *Dnmt1* in *β*-cells results in demethylation and depression of *Arx* ^40^. We thus speculate that *ARX* is hypomethylated in pancreatic progenitors in a TET-independent manner. The *ARX* locus maintains an unmethylated state when progenitors differentiate into *α*-cells, whereas it becomes hypermethylated once cells commit to *β*-cell fate in the presence of DNMTs and other *β* cell-specific regulators. In line with this hypothesis are findings that Nkx2.2 recruits Dnmt3a to the *Arx* promoter to repress its expression in β-cells ^42^. In the future, it will be interesting to explore whether there are differences in the complete epigenetic landscapes of endocrine progenitors, which subsequently differentiate into *α*- or *β*-cells.

TET dioxygenases are critical for lineage induction in a cell-type-specific manner ^43^. How TETs recognize lineage-specific regulatory elements and modulate chromatin remodeling during pancreas development remains unknown. Here, we addressed these questions by performing analyses integrating gene expression with multiple chromatin features, including DNA methylation, hydroxymethylation, chromatin accessibility, and histone modifications of enhancers. We found extensive hypermethylation in TET-deficient cells that differentiated to the pancreatic progenitor stage. A significant portion of hyper-DMRs was distributed in a differentiation-specific manner, in which they were enriched for the binding of FOXA2, a pioneer TF essential for pancreatic endoderm differentiation, and showed remarkable decreases in chromatin activity upon TET inactivation. It is noteworthy that we unveiled a unique TF binding pattern associated with a stepwise increase of 5hmC during pancreatic differentiation. The presence of GATA, FOXA, and HNF6 binding motifs in a temporal manner suggests that TET proteins are recruited to chromatin by a set of lineage-specific TFs sequentially expressed during pancreatic differentiation.

During lineage specification, chromatin structure is dynamically changed between ‘closed’ and ‘open’ states. Open chromatin regions such as primed and active enhancers are accessible for TF and epigenetic modulator binding to initiate gene transcription. Proper chromatin remodeling is believed to be at least partly triggered by pioneer TFs, which can directly bind to nucleosomal DNA ^44^. It was previously suggested that pioneer TF FOXA2 initiates chromatin remodeling and enhancer priming during pancreatic differentiation ^3, 4^. Interestingly, our analyses revealed that 1) changes in 5hmC mirror the dynamic binding of FOXA2 in cells differentiated from hESCs through defined lineage intermediates toward pancreatic endocrine fate, 2) chromatin activity was markedly decreased at the FOXA2 binding sites associated with hyper-DMRs in TET-deficient cells, and 3) FOXA2 physically interacted with TET1, suggesting that FOXA2 recruits TET1 mediate lineage-specific demethylation at a subset of FOXA2 targets to influence chromatin opening.

How pioneer TFs selectively recognize differentiation stage-specific target sites and mediate chromatin remodeling is still mostly unknown. Our present study demonstrates that 1) de novo FOXA2 recruitment to genomic loci of low levels of active chromatin modifications is inhibited upon TET depletion, and 2) FOXA2-decreased binding sites are enriched with bHLH TFs, such as NEUROD1 and PTF1A, which fail to be induced in TET-deficient cells. Interestingly, FOXA2 stable and continuous binding targets are largely absent of motifs except for the forkhead and CTCF family motifs. Therefore, we speculate that FOXA2 transient binding is facilitated by lineage-specific TFs in a differentiation-stage specific manner during pancreatic cell fate commitment (Fig. 6f). Indeed, it was recently demonstrated that chromatin opening requires cooperative binding of FOXA2 and additional TFs in close vicinity ^32^. However, why co-binding of TFs is necessary for initiating chromatin remodeling is unclear. Our findings suggest that de novo binding of FOXA2 to lineage-appropriate sites may require by cell-type-specific TFs, where chromatin modifiers such as TET1 are recruited to mediate subsequent chromatin opening. Further investigation is warranted to determine whether and how lineage-specific TFs facilitate FOXA2 binding and synergize the recruitment of multiple chromatin remodeling machineries.

## Methods

### Cell lines

The human embryonic stem cell line H1 was obtained from WiCell Research Institute. H1 cells were maintained on Matrigel (Corning) in mTeSR1 (STEMCELL Technologies) and passaged every for 4-5 days using 0.5 mM EDTA and 3 mM NaCl. Mycoplasma detection was regularly performed by the Stem Cell Core Facility at the University of Macau.

### Pancreatic differentiation

All differentiation experiments were repeated at least three times with three individual clones of the same phenotype. Pancreatic differentiation was performed as previously described with slight modifications ^45^. In brief, hESCs were dissociated with Accutase (eBioscience) and seeded at a density of 19,000 cells/cm^2^ on Matrigel in mTeSR1 supplemented with 10 μM Rho-associated protein kinase inhibitor Y-27632 (Miltenyi Biotec). Upon reaching 95% confluence, cells were exposed to differentiation medium with daily media feeding. Alternatively, a suspension-based differentiation format was used as previously described ^27^. Briefly, 5.5 × 10^6^ cells were suspended in 5.5 ml mTeSR1 per well of 6-well ultra-low attachment plates (Coring Costar) and cultured overnight in an orbital shaking CO_2_ incubator (Eppendorf New Brunswick) at 95 rpm. Cell aggregates were supplied with differentiation medium and continually rotated at 95 rpm with daily media feeding.

ES-to-DE (3 d): hESCs were exposed to 100 ng/ml Activin A (PeproTech) and 25 ng/ml Wnt-3a (PeproTech) in basal medium-I containing MCDB 131 medium (Gibco), 1.5 g/l NaHCO_3_, 1× GlutaMAX (Gibco), 10 mM glucose, and 0.5% BSA for 1 day. For 2 additional days, cells were cultured in basal medium-I further supplemented with 100 ng/ml Activin A.

DE-to-GT (3 d): DE intermediates were incubated in basal medium-I supplemented with 50 ng/ml FGF7 (PeproTech) for 3 days.

GT-to-PP (5 d): GT intermediates were cultured for 3 days in basal medium-II containing MCDB 131 medium, 2.5 g/l NaHCO_3_, 1× GlutaMAX, 10 mM glucose, 2% BSA, and 1:200 ITS-X (Gibco), which was further supplemented with 50 ng/ml FGF7, 0.25 μM hedgehog inhibitor SANT-1 (Sigma), 1 μM retinoic acid (Sigma), 100 nM BMP inhibitor LDN193189 (Stemgent), and 200 nM PKC activator TPB (Millipore). After 3 days of culture, cells were treated for 2 days with 2 ng/ml FGF7, 0.25 μM SANT-1, 0.1 μM retinoic acid, 200 nM LDN193189, and 100 nM TPB in basal medium-II.

PP-to-PE (3 d): PP intermediates were differentiated in basal medium-III containing MCDB 131 medium, 1.5 g/l NaHCO_3_, 1× GlutaMAX, 20 mM glucose, 2% BSA, and 1:200 ITS-X, which was further supplemented with 0.25 μM SANT-1, 0.05 μM retinoic acid, 100 nM LDN193189, 1 μM T3 (3,3’,5-Triiodo-L-thyronine sodium salt, Sigma), 10 μM ALK5 inhibitor II (Enzo Life Sciences), 10 μM ZnSO_4_, and 10 μg/ml heparin (Sigma).

### Animal experiments

All animal experiments were approved by the University of Macau Animal Ethics Committee. Immunocompromised SCID-beige mice were obtained from Charles River and maintained under a 12-h light/dark cycle with free access to standard mouse diet. For transplantation, d14 cell aggregates were further incubated in basal medium-III supplemented with 100 nM LDN193189, 1 μM T3, 10 μM ALK5 inhibitor II, 10 μM ZnSO_4_, and 100 nM *γ*-secretase inhibitor XX (Millipore) for 1 day. Cell aggregates (∼5 × 10^6^ cells) were then transplanted into 6- week-old SCID-beige mice under the kidney capsule as previously described ^46^.

Glucose-stimulated human C-peptide secretion was assessed with mice 18 weeks post-transplantation. Blood samples were collected after overnight fasting and 30 min following an intraperitoneal injection of glucose (2 g/kg body weight). Human C-peptide levels in isolated plasma were quantified using the STELLUX Chemi Human C-peptide ELISA kit (ALPCO Diagnostics) according to the manufacturer’s instructions.

### Generation of *TET* knock-out lines

*TET* knock-out hESCs were generated using CRISPR/Cas9 technology. gRNAs (Supplementary Table 5) were designed to target the sequences encoding exon 7 of *TET1*, exon 3 of *TET2*, or exon 3 of *TET3* and cloned into pX330 vector (Addgene #42230) as previously described ^47^. Constructs containing validated gRNAs were electroporated together with a vector expressing puromycin into hESCs using the P3 Primary Cell 4D-Nucleofector X kit (Lonza) following the manufacturer’s instructions. The electroporated cells were plated at ∼2,000 cells/cm^2^ and cultured on Matrigel in mTeSR1 supplemented with 10 μM Y-27632 for 2 days. Successfully transfected cells were selected with 1 μg/ml puromycin in mTeSR^TM^1 and allowed to expand to form visible colonies from a single cell. Subsequently, clonal colonies were manually picked and reseeded individually into 24-well plates. The amplified colonies were analyzed by Sanger sequencing at targeted loci for the presence of Indel mutations.

### TET1 overexpression cell lines

To generate overexpression constructs for TET1, the C-terminal FLAG-tagged human full-length TET1 (TET1FL), TET1 catalytic domain (TET1CD), and TET1 catalytic inactive mutant (TET1CD^mut^) were amplified from FH-TET1-pEF (Addgene #49792) or pIRES-hrGFP II-mTET1 (Addgene #83569), and cloned into the *NotI* digested PCDH-CAG-MCS-P2A-Puro vector (a kind gift from Dr. R. Xu, University of Macau) ^48^. Lentivirus particles were prepared as previously described ^49^. For viral transduction, TKO-hESCs were grown on Matrigel in mTeSR1 using 24-well plates and treated with 6 μg/ml polybrene for 15 min at 37°C. Concentrated lentivirus particles (10 μl, 1 × 10^6^ TU/ml) were added to cell culture and incubated overnight at 37°C. On the next day, viral infection was repeated (30 μl, 1 × 10^6^ TU/ml) to increase transduction efficiency. Infected cells were cultured in mTeSR1 for 2 days and then exposed to 1 μg/ml puromycin for 10 days. The TET1 overexpression cell lines TKO-TET1FL, TKO-TET1CD, TKO-TET1CD^mut^, were amplified and frozen down.

### Co-immunoprecipitation

Cells were lysed in ice-cold lysis buffer (20 mM Tris pH 7.5, 150 mM NaCl, 1 mM EDTA, 1% Triton X-100, 5mM 4-nitrophenyl phosphate di(Tris) salt, 2mM Na2VO4, 0.5% sodium deoxylcholate, protease inhibitors), sonicated on Diagenode Bioruptor for 5 min (30 s ON and 30 s OFF), and then centrifuged at 12,000 rpm for 10 min at 4 °C. Supernatants were incubated with pre-balanced ANTI-FLAG® M2 Affinity Gel (Sigma-Aldrich, A2220) overnight at 4 °C. The beads were washed three times with wash buffer I (20 mM Tris pH 8.0, 0.3 M KCl, 10% glycerol, 1 mM EDTA, 1 mM DTT, 0.1% NP-40 and proteinase inhibitors) and two times with wash buffer II (20 mM Tris pH 8.0, 0.1 M KCl, 10% glycerol, 1 mM EDTA, 1 mM DTT and proteinase inhibitors). 4 × Laemmli sample buffer with β-mercaptoethanol was added to washed beads, and then boiled for 10 min at 95 °C for western blotting analyses.

### Locus-specific detection of 5mC

Detection of 5mC content at particular CCGG sites was performed using the Epimark 5hmC and 5mC analysis kit (New England Biolabs) following the manufacturer’s instructions. Briefly, genomic DNA was extracted using the DNeasy Blood and Tissue kit (Qiagen) followed by RNase treatment. RNase-treated DNA samples were incubated with T4 β-glucosyltransferase at 37°C for 16 h. The glycosylated DNA was subsequently digested with *MspI* or *HpaII* for 8 h at 37°C. Samples were treated with Proteinase K at 40°C for 30 min and then at 95°C for 10 min to inactivate the enzymes. Site-specific methylation contents were examined by RT-qPCR using the primers listed in Supplementary Table 6. The percentage of 5mC and unmodified cytosine were calculated using the comparative C_t_ method.

### 5hmC dot blot assay

Genomic DNA was extracted using the DNeasy Blood and Tissue kit (Qiagen) according to the manufacturer’s instructions. DNA was denatured in 1 M NaOH supplemented with 25 mM EDTA at 95°C for 10 min and then neutralized with 2 M ice-cold ammonium acetate for 10 min. Two-fold serial dilutions of the DNA samples were spotted onto nitrocellulose membrane. The air-dried membrane was fixed with UV irradiation (CL-1000 UV crosslinker, Ultra-Violet Products), blocked with 5% non-fat milk, and incubated with a rabbit anti-5hmC antibody (1:10,000, Active Motif) followed by an HRP-conjugated anti-rabbit antibody (1:5,000, Jackson ImmunoResearch). Signal was visualized with SuperSignal West Pico PLUS chemiluminescent substrate (Thermo Fisher Scientific). The same membrane was subsequently stained with 0.02% methylene blue in 0.3 M sodium acetate to ensure equal loading of input DNA.

### RNA isolation for real-time quantitative PCR

Total RNA was extracted using the RNeasy Plus Mini kit (Qiagen) according to the manufacturer’s instructions. cDNA was synthesized using the PrimeScript RT reagent kit (Takara). Real-time quantitative PCR was performed in triplicate using the SYBR Premix Ex Taq (Tli RNase H Plus) kit (Takara). The expression of *TBP* was used for the normalization of mRNA expression. All primers used for RT-qPCR were listed in Supplementary Table 6.

### Immunocytochemical analysis

Differentiated cells were fixed in 4% paraformaldehyde for 30 min at room temperature, washed, and then permeabilized with 0.15% Triton X-100 at room temperature for 1 h. Following blocking with 5% normal donkey serum (Jackson Immuno Research Laboratories), samples were incubated with primary antibodies at 4°C overnight and then appropriate secondary antibodies for 1 h at room temperature. Images were acquired using the Zeiss Axio Observer microscope. For transplant grafts, tissues were fixed with 4% paraformaldehyde overnight at 4°C, washed with PBS, and subsequently exposed to 30% sucrose overnight at 4°C. Samples were mounted with Optimal Cutting Temperature Compound (Tissue-Tek) and sectioned at 10 µm. Immunofluorescent staining was performed on cryosections as described above. Antibodies used for immunofluorescent staining are listed in Supplementary Table 7.

### Flow cytometry analysis

Cells were enzymatically dissociated into single cells and fixed with 4% paraformaldehyde for 20 min at 4°C. Fixed cells were permeabilized with Perm/Wash Buffer (BD Biosciences) and stained with primary antibodies (Supplementary Table 7) diluted in Perm/Wash Buffer overnight at 4°C. Cells were subsequently washed, stained with appropriate secondary antibodies for 1 h at room temperature, and assessed using an Accuri C6 flow cytometer (BD Biosciences). Data were analyzed using FlowJo software (TreeStar).

### RNA-seq library construction and data analysis

The integrity of extracted RNA was determined using an Agilent 2100 Bioanalyzer (Agilent Technologies). Subsequently, polyA-tailed RNA was selected using Dynabeads oligo (dT) (Thermo Fisher Scientific), and libraries were prepared using the NEBNext Ultra RNA Library Prep kit for Illumina (New England Biolabs). Libraries were subjected to high-throughput sequencing on an Illumina HiSeq 2500 system (150 cycles, paired-end) at Novogene (Tianjin, China). To process the sequencing data, low-quality bases and adaptor were trimmed using TrimGalore v.0.5.0 (options: --quality 20 and --length 50, https://github.com/FelixKrueger/TrimGalore). Clean reads were aligned to the hg38 genome reference using STAR v.2.5.3 ^50^ with default parameters, and only uniquely mapped reads were used for downstream analysis. A count matrix for each gene was generated using htseq-count (HTSeq package ^51^). DESeq2 ^52^ was used to identify significant DEGs in knock-out samples compared with WT samples at different differentiation stages (|fold change| ≥ 2; False Discovery Rate (FDR) < 0.05). Hierarchical cluster analysis of the union DEGs was used to determine stage-specific signature genes. The ‘ClusterProfilter’ package in R was used for the functional enrichment analysis of DEGs in KEGG pathways. BAM files were converted to bigwig files by Bam2wig.py in RSeQC ^53–55^.

### ATAC-seq library preparation and data analysis

ATAC-seq libraries were prepared as previously described ^56^. In brief, cells were enzymatically dissociated and lysed in lysis buffer (10 mM Tris-HCl, pH 7.4, 10 mM NaCl, 3 mM MgCl2, 0.1% IGEPAL CA-630). Immediately after centrifugation, transposition reactions were carried out by adding Tn5 transposes from the Illumina Nextera DNA library preparation kit to the isolated nuclei and incubation at 37°C for 30 min. DNA fragments were purified using the MinElute PCR Purification kit (Qiagen) and amplified using the KAPA real-time library amplification kit (Roche). Libraries were purified using PCRClean DX beads (Aline Biosciences) and subsequently subjected to high-throughput sequencing on an Illumina NextSeq 500 instrument (75 cycles, paired-end).

Adaptor trimming of raw reads was performed using TrimGalore v0.5.0 (options: --quality 20 and --length 20), and high-quality (Q ≥ 20) reads were uniquely aligned to the hg38 genome reference using Bowtie2 with the ‘--very-sensitive’ option. Reads mapped to mitochondrial DNA and PCR duplicate reads were removed, and uniquely mapped reads were extracted for downstream analysis. Genrich v.0.5 (https://github.com/jsh58/Genrich) with ATAC-seq mode (option: -j, -q 0.01) was applied for each sample (with two biological replicates) to call ATAC-seq peaks. In total, 52,817 and 38,697 peaks were detected in WT_PP and TKO_PP cells, respectively. BEDTools ^57^ intersect was used to generate a total of 124,322 non-overlapping peak regions, and DeepTools multiBamSummary was applied to count the reads falling into peak regions. The counts were used in DESeq2 for normalization and identification of significant differential accessible regions (DARs) between WT_PP and TKO_PP cells by the criteria (|Fold change| ≥ 2; FDR < 0.05). Volcano plots were generated using the R package ggplot2 ^58^. Motif annotation of DARs was performed using HOMER software ^59^.

### CMS-IP-seq library preparation and data analysis

CMS-IP-seq libraries were performed as previously described with minor modifications (Huang et al., 2012). Purified genomic DNA (with 5% mouse DNA and 0.5% lambda DNA spike-in) was sheared to 200 - 500 bp fragments using a M220 Focused-ultrasonicator (Covaris). Bisulfite-converted DNA libraries with methylated adapters were enriched using an in-house anti-CMS antibody bound to protein A/G Dynabeads. Amplified libraries were purified by AmpuXP beads (Beckman Coulter) and then sequenced using the Illumina NextSeq (75 and 40 cycles, single-end) system.

Analysis of CMS-IP-seq data was performed by in-house software ‘HaMiP’ v0.1.1 (https://github.com/lijinbio/HaMiP). Briefly, raw reads were mapped to hg38 and spike-in mm10 genome references using bsmap v.2.89 (options: -n 1 -q 3 -r 0) ^60^. After removing PCR duplicates and reads mapped to both human and spike-in mouse genome, species-specific reads were used to perform normalization for each sample according to the spike-in size factors. The whole human genome was divided into 200-bp windows and the normalized mean wigsum in each window was calculated to call the hydroxymethylation-enriched peak regions for each sample against input control. In total, 75,324 and 503 peaks were detected in WT_PP and TKO_PP samples, respectively. Differentially DHMRs between WT_PP and TKO_PP were identified (*G*-test; FDR < 0.05). GREAT analysis with single-nearest genes option was used to perform functional annotation of hypo-DHMRs.

CMS-IP-seq datasets of hESCs and multiple pancreatic lineage intermediates (DE, GT, and PP) were downloaded from GSE97992 ^20^ and mapped to the hg38 genome reference using bsmap with the same parameters as in the previous analysis. Bam2wig.py was used to transform the BAM file to normalized bigwig files (option: -t 2000000000). The average 5hmC signals within each hypo-DHMR for all stages were calculated using DeepTools ^61^. Compared with hESCs, increased 5hmC peaks were defined as peaks in PP cells with a ≥ 1.5-fold increase.

### WGBS library preparation and data analysis

Genomic DNA was isolated using the DNeasy Blood & Tissue kit (Qiagen). Library preparation and high-throughput sequencing were conducted by BGI (Shengzhen, China). In brief, purified genomic DNA (with 1% unmethylated lambda DNA spike-in, Promega) was sheared to a fragment size of 100 - 700 bp (primary size 250 bp). Sheared DNA was ligated with methylated adaptors (MGIEasy WGBS Adapters-16, MGI) and subjected to bisulfite conversion using the EZ DNA Methylation-gold kit (Zymo Research). Bisulfite-converted DNA was amplified with 13 PCR cycles and purified by AMPure XP beads (Beckman Coulter). All libraries were sequenced on an MGISEQ-2000 system (100 cycles, paired-end).

For data analysis, raw reads were mapped against hg38 genome reference using bsmap v.2.89 ^60^ with paired mode (options: -n 1 -q 3 -r 0), and only uniquely mapped reads were retained. More than 26.6 million CpG sites with coverage of ≥ 5 reads were detected in both WT_PP and TKO_PP samples, which were used for downstream analyses. BSeQC ^62^ and the Mcall module in MOABS ^28^ was applied to perform quality control and calculate the methylation ratio for each CpG site (options: --trimWGBSEndRepairPE2Seq 40). Bisulfite conversion efficiencies were estimated using spike-in unmethylated lambda phage DNA. The Mcomp module was used to call significant differentially methylated CpG sites and regions (DMCs and DMRs, absolute credible difference of DNA methylation ratio > 20% and FDR < 0.05). WGBS data of hESC was downloaded from the ENCODE database ^29^, and the average 5mC level of each hyper-DMR in hESC was also estimated. Detected hyper-DMRs (up- and down- 1 kb) were divided into 10-bp bins, and the average methylation level of each bin was calculated for hESC, WT_PP and TKO_PP samples, independently. The output bedGraph files from Mcall included single-base resolution DNA methylation ratios, which were transformed into bigwig file format. Motif enrichment analysis of DMRs was performed using HOMER software, functional annotation was performed using GREAT with default settings, and many plots related to WGBS data was performed using in-house software Mmint (https://github.com/lijiacd985/Mmint).

### ChIP-seq library preparation and data analysis

ChIP-seq was performed for FOXA2 (WT_DE, WT_GT, WT_PP, TKO_DE, and TKO_PP cells), H3K4me1 (WT_PP and TKO_PP cells), and H3K27ac (WT_PP and TKO_PP cells). Chromatin immunoprecipitation was performed as previously described ^27^. In brief, ∼1 × 10^7^ cells were crosslinked, washed, and lysed in nuclear lysis buffer (50 mM Trish HCl, pH 8.0, 5 mM EDTA, 1% SDS, and 1*×* protease inhibitor cocktail). Chromatin was sheared to 200–500-bp fragments using Bioruptor (Diagenode), and DNA fragments were precipitated with appropriate antibodies. ChIP-seq libraries were prepared using the NEB Next Ultra II DNA Library Prep kit (New England Biolabs) following the manufacturer’s instruction and subjected to high-throughput sequencing on an Illumina HiSeq 2500 system (150 cycles, paired-end) at Novogene (Tianjin, China).

Quality control and alignment of raw reads of ChIP-seq data were performed similarly to the ATAC-seq data analysis described above. TrimGalore (options: --quality 20 and --length 50) was used to remove the adaptor. Bowtie2 with the ‘--very-sensitive’ option was used for alignment, and only uniquely mapped reads were retained. Bam2wig.py was used to transform the BAM file to normalized bigwig files for visualization (option: -t 2000000000). For FOXA2 ChIP-seq, Macs2 ^63^ was used to call ChIP-seq enriched peak regions with default parameters for each replicate. The BEDTools intersect was used to obtain the highly confident FOXA2 bound regions in two biological replicates for each sample. Then BEDTools merge was used to merge the confident peaks from WT and TKO samples to create the consensus FOXA2 bound regions at the DE and PP stage, independently. The reads number was counted for consensus peaks in each replicate and DESeq2 was used to call significant differential FOXA2 bound regions (FC≥ 2, FDR < 0.05). Similar to ATAC-seq, motif analysis of FOXA2 bound regions was performed using HOMER software and functional annotation was performed using GREAT.

### Integration of analyses

RNA-seq, ATAC-seq, WGBS, CMS-IP-seq and ChIP-seq library preparations were performed as previously described. Detail experimental procedures can be found in Supplementary Materials. The number of mapped reads, mapped ratios and other statistic information are listed in Supplementary Table 8. To compare DNA methylation between TKO_PP and WT_PP cells at annotated genomic features, ChIP-seq datasets of GATA4, GATA6, and PDX1 were downloaded from GSE117136. Bivalent promoters and poised and active enhancers in hESC-derived pancreatic progenitors were downloaded from EMTAB1086 and GSE54471, respectively. Data were analyzed as described above. The methylation difference between WT_PP and TKO_PP across each genomic feature was performed by deepTools and Mmint software.

For integrated data visualization, bigwig files from RNA-seq, ATAC-seq, WGBS, CMS-IP-seq and ChIP-seq experiments were uploaded to the University of California, Santa Cruz (UCSC) genome browser. R package ggplot2 was used to plot volcano, bar, boxplot, point, heatmap and scatter plots ^58^.

### Quantification and statistical analysis

All statistical analyses were performed using GraphPad Prism software. Student’s unpaired two-tailed *t*-tests were used for qRT-PCR and flow cytometry experiments. Student’s paired one-tailed *t*-tests were used for the glucose-stimulated human C-peptide secretion experiment. Quantification data are presented as mean ± SD. For all statistical analyses, *p < 0.05, **p < 0.01, ***p < 0.001, and ****p < 0.0001. Statistical analyses for RNA-seq, WGBS, CMS-IP-seq, and ChIP-seq data are described in the corresponding sections.

### Data availability

WGBS, CMS-IP, RNA-seq, ATAC-seq, and ChIP-seq data from this study were submitted to the NCBI Gene Expression Omnibus (GEO; https://www.ncbi.nlm.nih.gov/geo/) under accession number GSE146486. All relevant data supporting the critical findings of this study are available within the article and its supplementary information files or from the corresponding author upon reasonable request.

## Acknowledgments

We thank the University of Macau, Faculty of Health Sciences, Animal Research Facility for animal housing. This work was supported by National Natural Science Foundation of China (NSFC 31701276) and The Science and Technology Development Fund, Macau SAR (0022/2019/AMJ) to R.X. The authors also thank the members of Macau Society for Stem Cell Research (MSSCR) for inspiring discussion.

## Author contributions

R.X. conceived the project. R.X. and D.S. directed and oversaw the project. X.W. and J. K. performed experiments and collected data. J.F.L. performed computational analysis for WGBS, RNA-seq, CMS-IP-seq, ATAC-seq, and ChIP-seq. M.L. prepared the CMS-IP library and performed sequencing. Q.L. assisted with the animal studies. J.L. provided expert advice on bioinformatic analyses. Y.H. critically reviewed the manuscript. R.X., J.F.L., X.W., and D.S. wrote the manuscript; all other authors provided editorial advice.

---

**Supplementary Fig. 1.**
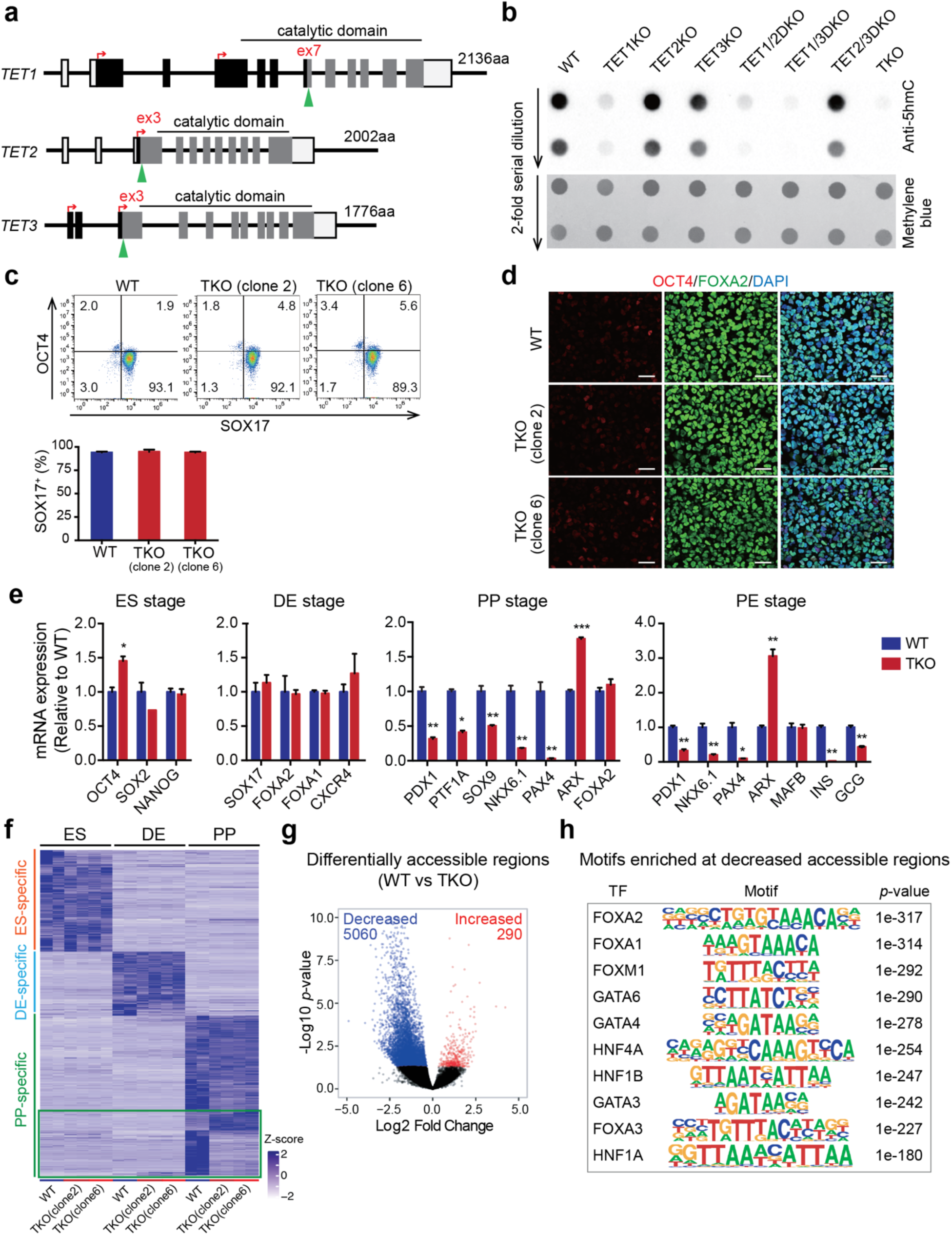
Generation and characterization of TET-deficient cell lines. **a** TET knock-out mutants were generated using CRISPR gRNAs (green arrowheads) targeting exon 7 of *TET1,* exon 3 of *TET2*, and exon 3 of *TET3*. Translation start sites are indicated by red arrows. **b** Analysis of global 5-hydroxymethylcytosine (5hmC) levels (top) in WT, *TET1* knock-out (TET1KO), *TET2* knock-out (TET2KO), *TET3* knock-out (TET3KO), *TET1/TET2* double knock-out (TET1/2DKO), *TET1/TET3* double knock-out (TET1/3DKO), *TET2/TET3* double knock-out (TET2/3DKO), and *TET1/TET2/TET3* triple knock-out (TKO) hESCs by 5hmC dot blot analysis. The bottom panel shows methylene blue staining using the total amount of input DNA as the loading control. **c** Representative plots of flow cytometry for the expression of pluripotency marker (OCT4) and endoderm marker (SOX17) at the DE stage are shown in the top panel. Quantification of the percentage of SOX17^+^ cells is shown in the bottom panel. n = 3 independent differentiation. **d** Immunostaining of OCT4 and FOXA2 in WT and TKO cells at the DE stage. Scale bar = 50 μm. **e** Expression analysis by RT-qPCR for specific genes in WT and TKO cells at the ES, DE, PP, and PE stages. RT-qPCR validation was performed with three independent batches of samples. **f** Heatmap showing stage-specific DEGs in TKO compared with WT cells at the ES, DE, and PP stages (|fold change| ≥ 2; FDR < 0.05). Two mutant lines were used for TKO (clones 2 and 6). Each column represents one biological replicate for each cell line. Genes located in green rectangle are DEGs in TKO_PP compared with WT_PP cells. **g** Volcano plot of ATAC-seq data illustrating differentially accessible regions in TKO_PP cells (FDR < 0.05). **h** Transcription factor (TF) motif enrichment analysis of genomic regions showed significantly decreased ATAC-seq signals in TKO_PP cells. The top 10 significant motifs are shown after removing redundant motifs.

**Supplementary Fig. 2.**
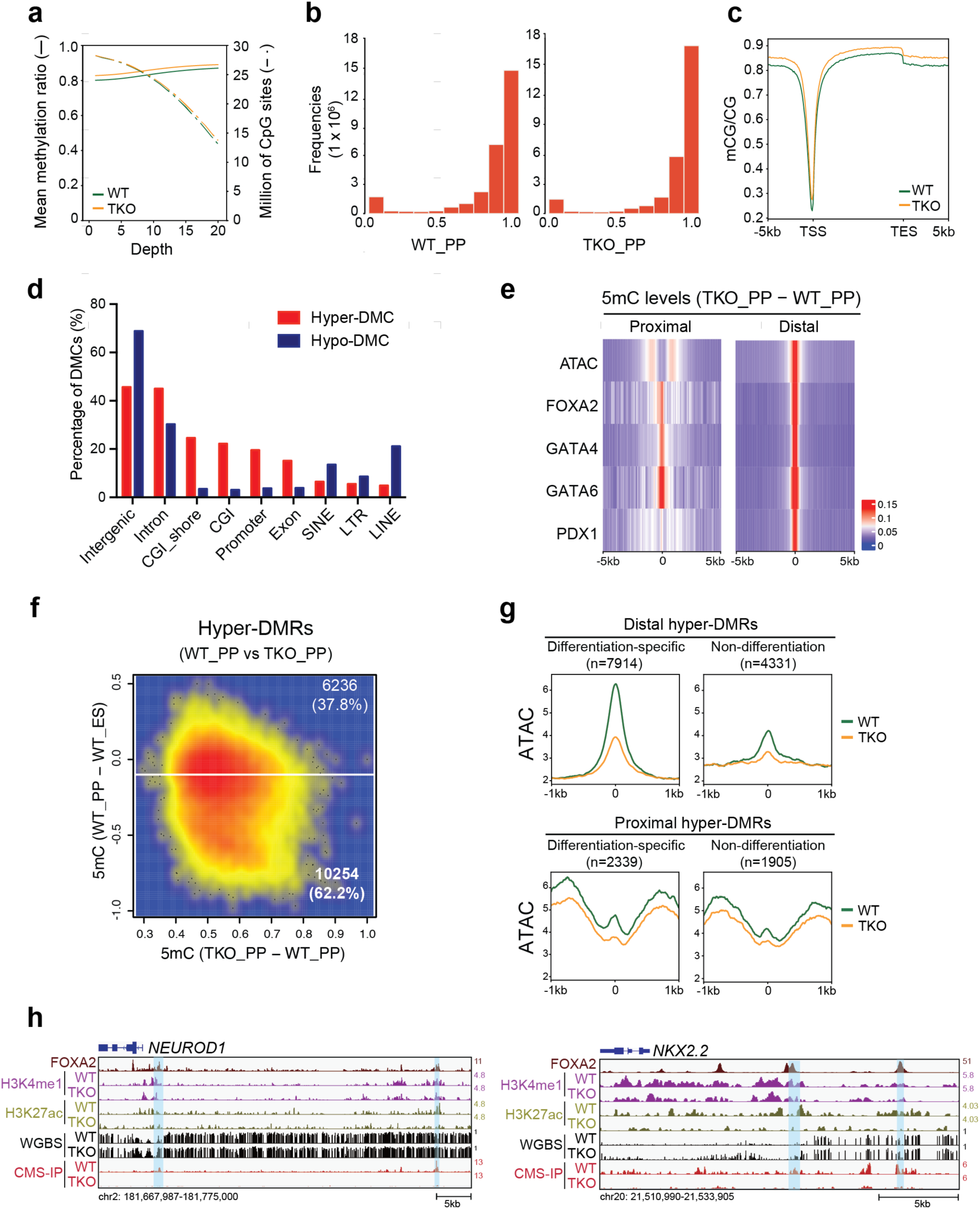
Loss of TET leads to extensive hypermethylation of pancreatic regulatory elements. **a** CpG coverage in WT_PP and TKO_PP cells. The x-axis represents the sequencing coverage for CpG sites. The solid line represents the average methylation ratio (5mC/C) at the corresponding coverage for each sample (left y-axis). The dashed line represents the number of CpGs over the corresponding coverage for each sample (right y-axis). **b** Bar plots showing the distribution of methylation in WT_PP and TKO_PP cells. **c** Mean distribution of the average methylation ratio (5mC/C) across gene bodies in WT_PP and TKO_PP cells. **d** Distribution of hyper-DMCs (red) and hypo-DMCs (blue) at various genomic features. **e** Heatmap illustrating methylation difference between TKO_PP and WT_PP cells at centers of annotated genomic features (± 5 kb) for chromatin accessibility (ATAC) and TF binding (FOXA2, GATA4, GATA6, and PDX1) at proximal (≤ 1 kb from TSS) and distal (> 1 kb from TSS) regions. **f** Comparison of global methylation at hyper-DMRs revealed that TKO_PP cells show differentiation-specific hyper-DMRs (n = 10,253) with decreased methylation in pancreatic progenitors (WT_PP) compared with hESCs (WT_ES) (methylation change < -0.1). **g** Average density plots of ATAC-seq signals at distal and proximal differentiation-specific or non-differentiation hyper-DMRs in WT_PP (green) and TKO_PP (orange) cells. **h** Genome-browser views of *NEUROD1* and *NKX2.2* loci. Representative hyper-DMRs showing decreased 5hmC and H3K27ac signals are highlighted in blue.

**Supplementary Fig. 3.**
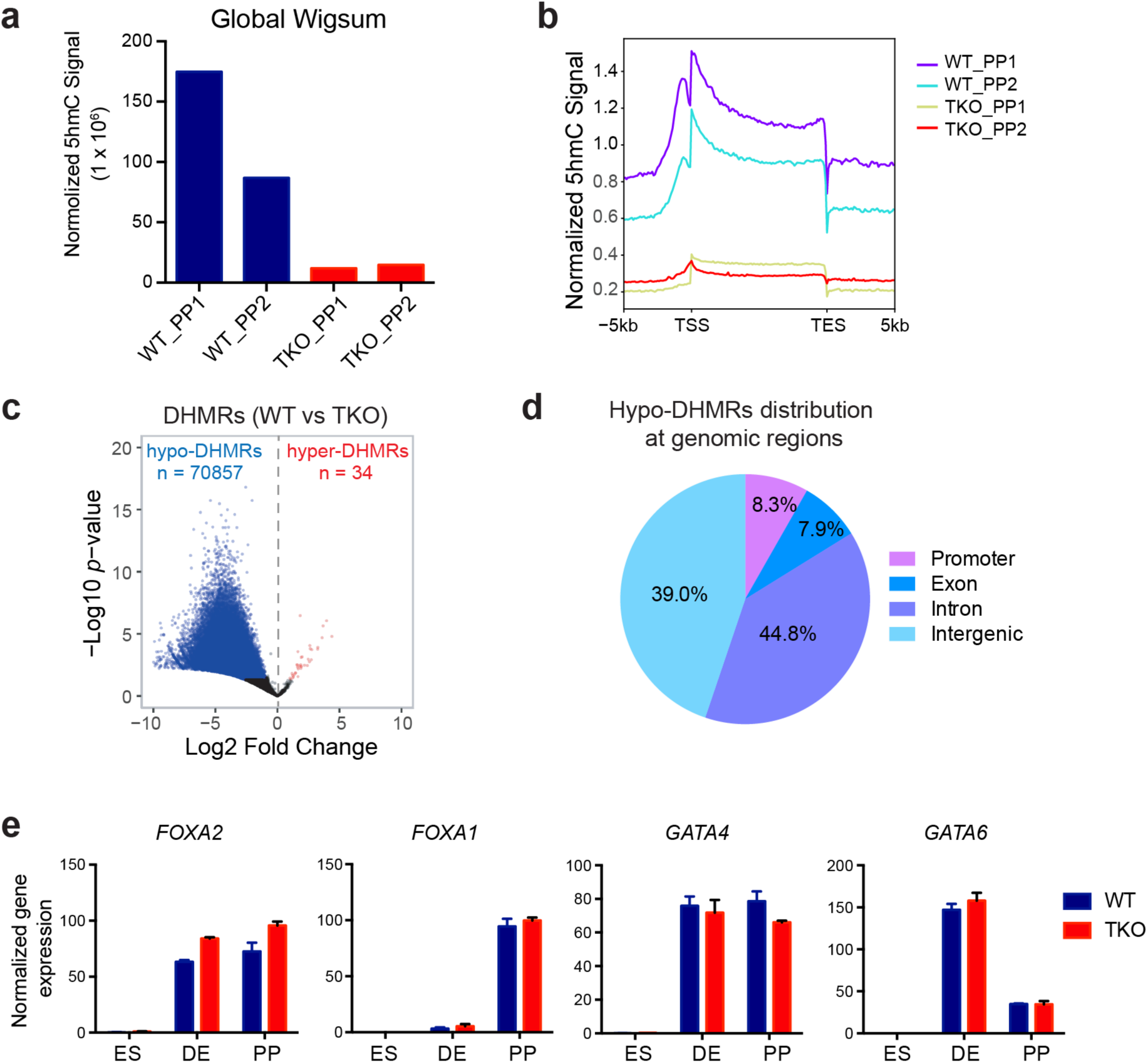
Hypo-hydroxymethylation at cis-regulatory elements upon TET inactivation. **a** Normalized 5hmC signal according to the spike-in analysis for WT_PP and TKO_PP samples with two independent replicates. **b** Distribution of 5hmC signal across gene bodies, from the transcription start site (TSS) to transcription end site (TES), in WT_PP and TKO_PP cells, with two independent replicates (± 5 kb). **c** Volcano plot of CMS-IP-seq signals illustrating differential hydroxymethylation regions (DHMRs) in TKO_PP cells relative to WT_PP cells (FDR < 0.05). **d** Diagram illustrating the overall distribution of hypo-DHMRs in promoter (within ± 1 kb from TSS), exon, intron, and intergenic regions. **e** Normalized gene expression of *FOXA1*, *FOXA2*, *GATA4*, and *GATA6* in WT and TKO cells at the ES, DE, and PP stages by RNA-seq.

**Supplementary Fig. 4.**
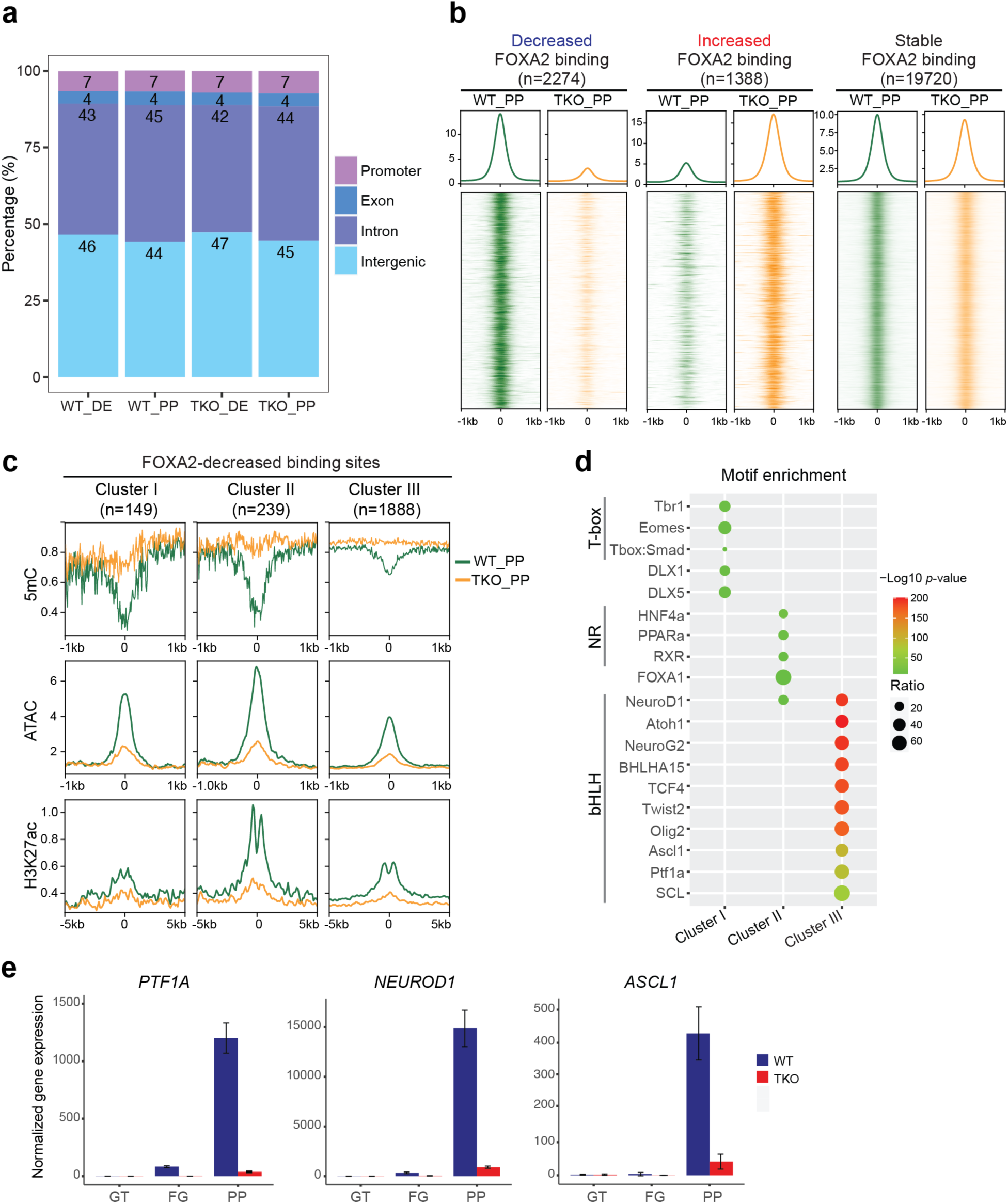
De novo FOXA2 binding at pancreas-specific loci features low active chromatin modifications. **a** Bar graph illustrating the percentage of FOXA2 binding sites associated with different genomic features in WT_DE, WT_PP, TKO_DE, and TKO_PP cells. **b** Average density plots and heatmaps of FOXA2 signals in WT_PP (green) and TKO_PP (orange) cells across decreased, increased, and stable FOXA2-bound sites. **c** Average density plot of methylation ratio (5mC/C), chromatin accessibility (ATAC), and H3K27ac at cluster III decreased FOXA2-bound regions in WT_PP (green) and TKO_PP (orange) cells. **d** Transcription factor motif enrichment analysis of genomic regions identified in cluster I, cluster II, and cluster III of decreased FOXA2 binding in TKO_PP cells. Only the top scoring motifs are shown. The color representing *p*-values and the size of circle representing proportion of peaks containing a TF motif in each group. **e** Normalized gene expression of *PTF1A*, *NEUROD1*, and *ASCL1* in WT and TKO cells at the GT, FG, and PP stages by RNA-seq.

**Supplementary Fig. 5.**
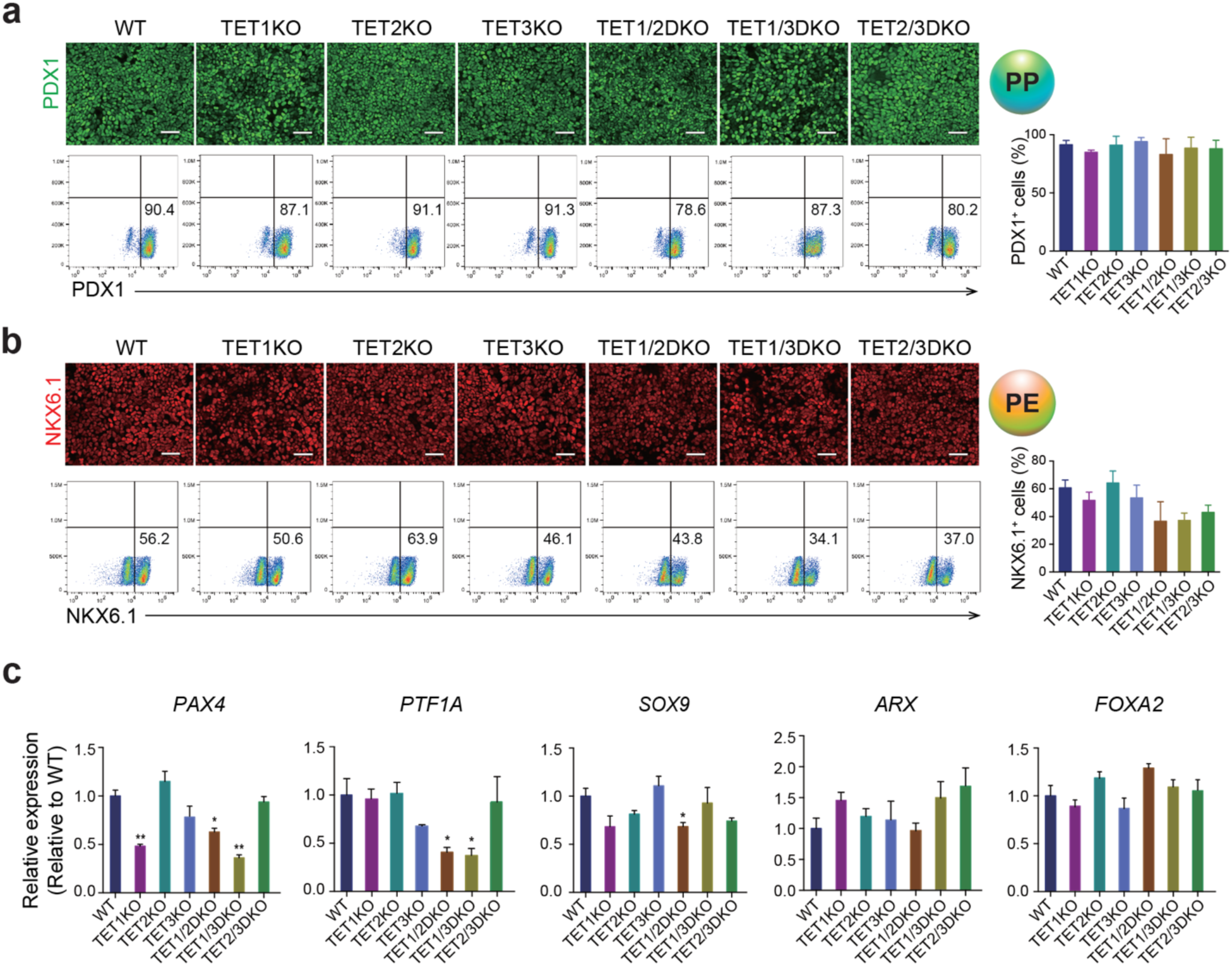
TET1 is required in pancreatic β-cell specification. **a** Immunostaining and representative plots of flow cytometry of PDX1 at the PP stage for WT, TET1KO, TET2KO, TET3KO, TET1/2DKO, TET1/3DKO, and TET2/3DKO cells. Quantifications of the percentage of PDX1^+^ cells are shown in the right panel. n = 3 independent differentiation. Scale bar = 50 μm. **b** Immunostaining and representative plots of flow cytometry of NKX6.1 at the PE stage for WT, TET1KO, TET2KO, TET3KO, TET1/2DKO, TET1/3DKO, and TET2/3DKO cells. Quantifications of the percentage of NKX6.1^+^ cells are shown in the right panel. n = 3 independent differentiation. Scale bar = 50 μm. **c** Expression of *PAX4*, *PTF1A*, *SOX9*, *ARX*, and *FOXA2* in WT, TET1KO, TET2KO, TET3KO, TET1/2DKO, TET1/3DKO, and TET2/3DKO cells at the PP stage by RT-qPCR. RT-qPCR validation was performed with three independent batches of samples.

**Supplementary Fig. 6.**
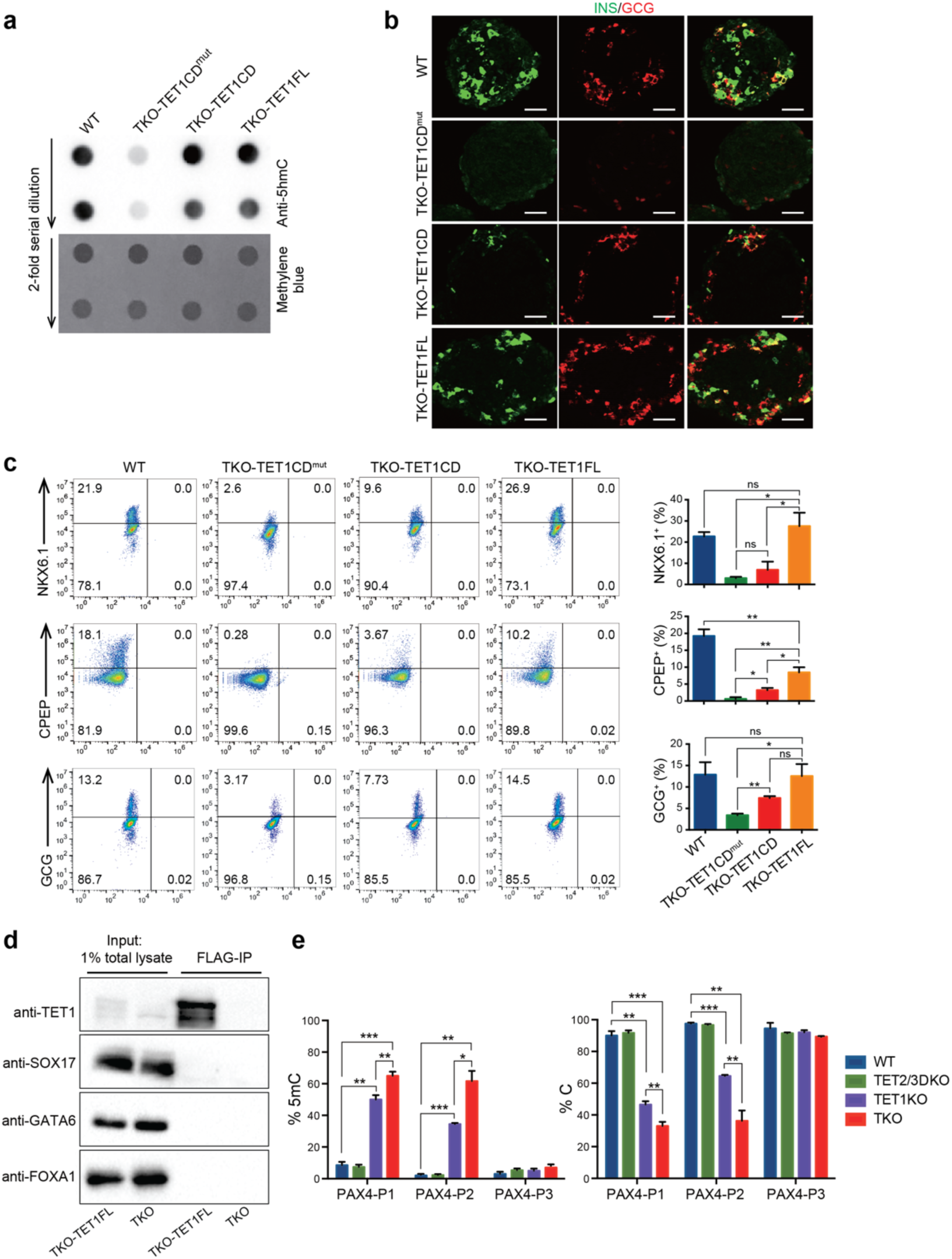
Reverse hypermethylation at *PAX4* enhancer through overexpression of full-length TET1. **a** Analysis of global 5-hydroxymethylcytosine (5hmC) levels (top) in WT, TKO-TET1CD^mut^, TKO-TET1CD, and TKO-TET1FL cells by 5hmC dot blot analysis. The bottom panel shows methylene blue staining using the total amount of input DNA as the loading control. **b** Immunostaining of insulin (INS) and glucagon (GCG) at the PE stage for WT, TKO-TET1CD^mut^, TKO-TET1CD, and TKO-TET1FL cells. Scale bar=50 μm. **c** Representative plots of flow cytometry of NKX6.1, human C-peptide (CPEP), and glucagon (GCG) in WT, TKO-TET1CD^mut^, TKO-TET1CD, and TKO-TET1FL cells at the PE stage. Quantifications of the percentage of NKX6.1^+^, CPEP^+^, or GCG^+^ cells are shown in the right panel. n = 3 independent differentiation. **d** Western blots showing immunoprecipitation with antibodies against FLAG on cell extract of TKO-TET1FL cells followed by immunoblotting with antibodies against TET1, SOX17, GATA6, or FOXA1. TKO is included as control. **e** Locus-specific decreases in 5-methylcytosine (5mC) at the *PAX4* enhancer in TKO or TET1KO samples compared with TET2/3DKO samples. Percentages of unmethylated cytosine and 5mC at CCGG sites are shown. n = 3 independent differentiation.

## References

1. Buecker, C. & Wysocka, J. Enhancers as information integration hubs in development: Lessons from genomics. Trends Genet. 28, 276–284 (2012).

2. Long, H. K., Prescott, S. L. & Wysocka, J. Ever-Changing Landscapes: Transcriptional Enhancers in Development and Evolution. Cell 167, 1170–1187 (2016).

3. Wang, A. et al. Epigenetic Priming of Enhancers Predicts Developmental Competence of hESC-Derived Endodermal Lineage Intermediates. Cell Stem Cell 16, 386–399 (2015).

4. Lee, K. et al. FOXA2 Is Required for Enhancer Priming during Pancreatic Differentiation. Cell Rep. 28, 382–393 (2019).

5. Cirillo, L. A. & Zaret, K. S. An Early Developmental Transcription Factor Complex that Is More Stable on Nucleosome Core Particles Than on Free DNA. Mol. Cell 4, 961–969 (1999).

6. Iwafuchi-Doi, M. et al. The Pioneer Transcription Factor FoxA Maintains an Accessible Nucleosome Configuration at Enhancers for Tissue-Specific Gene Activation. Mol. Cell 62, 79–91 (2016).

7. Donaghey, J. et al. Genetic determinants and epigenetic effects of pioneer-factor occupancy. Nat. Genet. 50, 250–258 (2018).

8. Geusz, R. J. et al. FoxA-dependent demethylation of DNA initiates epigenetic memory of cellular identity. bioRxiv (2020). doi:10.1016/j.devcel.2021.02.005

9. He, Y. F. et al. Tet-Mediated Formation of 5-Carboxylcytosine and Its Excision by TDG in Mammalian DNA. Science 333, 1303–1307 (2011).

10. Ito, S. et al. Role of tet proteins in 5mC to 5hmC conversion, ES-cell self-renewal and inner cell mass specification. Nature 466, 1129–1133 (2010).

11. Ito, S. et al. Tet proteins can convert 5-methylcytosine to 5-formylcytosine and 5-carboxylcytosine. Science 333, 1300–1303 (2011).

12. Tahiliani, M. et al. Conversion of 5-methylcytosine to 5-hydroxymethylcytosine in mammalian DNA by MLL partner TET1. Science 324, 930–935 (2009).

13. Zhang, H. et al. TET1 is a DNA-binding protein that modulates DNA methylation and gene transcription via hydroxylation of 5-methylcytosine. Cell Res. 20, 1390–1393 (2010).

14. Tsagaratou, A. et al. Dissecting the dynamic changes of 5-hydroxymethylcytosine in T-cell development and differentiation. Proc. Natl. Acad. Sci. USA 111, E3306–E3315 (2014).

15. Tan, L. et al. Genome-wide comparison of DNA hydroxymethylation in mouse embryonic stem cells and neural progenitor cells by a new comparative hMeDIP-seq method. Nucleic Acids Res. 41, e84 (2013).

16. Kim, M. et al. Dynamic changes in DNA methylation and hydroxymethylation when hES cells undergo differentiation toward a neuronal lineage. Hum. Mol. Genet. 23, 657–667 (2014).

17. Verma, N. et al. TET proteins safeguard bivalent promoters from de novo methylation in human embryonic stem cells. Nat. Genet. 50, 83–95 (2018).

18. Ko, M. et al. Ten-eleven-translocation 2 (TET2) negatively regulates homeostasis and differentiation of hematopoietic stem cells in mice. Proc. Natl. Acad. Sci. USA 108, 14566–14571 (2011).

19. Fang, S. et al. Tet inactivation disrupts YY1 binding and long-range chromatin interactions during embryonic heart development. Nat. Commun. 10, 1–18 (2019).

20. Li, J. et al. Decoding the dynamic DNA methylation and hydroxymethylation landscapes in endodermal lineage intermediates during pancreatic differentiation of hESC. Nucleic Acids Res. 46, 2883–2900 (2018).

21. Geusz, R. J. et al. A dual mechanism of enhancer activation by FOXA pioneer factors induces endodermal organ fates. bioRxiv (2020).

22. Lu, F., Liu, Y., Jiang, L., Yamaguchi, S. & Zhang, Y. Role of Tet proteins in enhancer activity and telomere elongation. Genes Dev. 28, 2103–2119 (2014).

23. Rodríguez-Seguí, S., Akerman, I. & Ferrer, J. GATA believe it: New essential regulators of pancreas development. J. Clin. Invest. 122, 3469–3471 (2012).

24. Kaestner, K. H. The FoxA factors in organogenesis and differentiation. Curr. Opin. Genet. Dev. 20, 527–532 (2010).

25. López-Moyado, I. F. et al. Paradoxical association of TET loss of function with genome-wide DNA hypomethylation. Proc. Natl. Acad. Sci. USA 116, 16933–16942 (2019).

26. Walsh, C. P., Chaillet, J. R. & Bestor, T. H. Transcription of IAP endogenous retroviruses is constrained by cytosine methylation. Nat. Genet. 20, 116–117 (1998).

27. Xie, R. et al. Dynamic chromatin remodeling mediated by polycomb proteins orchestrates pancreatic differentiation of human embryonic stem cells. Cell Stem Cell 12, 224–237 (2013).

28. Sun, D. et al. MOABS: model based analysis of bisulfite sequencing data. Genome Biol. 15, 1–12 (2014).

29. Consortium, E. P. An integrated encyclopedia of DNA elements in the human genome. Nature 489, 57–74 (2012).

30. Zhang, H. et al. Multiple, temporal-specific roles for HNF6 in pancreatic endocrine and ductal differentiation. Mech. Dev. 126, 958–973 (2009).

31. Li, Z. et al. Foxa2 and H2A.Z Mediate Nucleosome Depletion during Embryonic Stem Cell Differentiation. Cell 151, 1608–1616 (2012).

32. Cernilogar, F. M. et al. Pre-marked chromatin and transcription factor co-binding shape the pioneering activity of Foxa2. Nucleic Acids Res. 47, 9069–9086 (2019).

33. Naya, F. J. et al. Diabetes, defective pancreatic morphogenesis, and abnormal enteroendocrine differentiation in BETA2/NeuroD-deficient mice. Genes Dev. 11, 2323– 2334 (1997).

34. Sosa-pineda, B., Chowdhury, K., Torres, M., Oliver, G. & Gruss, P. The Pax4 gene is essential for differentiation of insulin-producing beta cells in the mammalian pancreas. Nature 386, 399–402 (1997).

35. Nelson, S. B., Schaffer, A. E. & Sander, M. The transcription factors Nkx6.1 and Nkx6.2 possess equivalent activities in promoting beta-cell fate specification in Pdx1+ pancreatic progenitor cells. Development 134, 2491–2500 (2007).

36. Ohta, Y. et al. Convergence of the insulin and serotonin programs in the pancreatic β-cell. Diabetes 60, 3208–3216 (2011).

37. Ahlgren, U., Jonsson, J., Jonsson, L., Simu, K. & Edlund, H. β-cell-specific inactivation of the mouse Ipf1/Pdx1 gene results in loss of the β-cell phenotype and maturity onset diabetes. Genes Dev. 12, 1763–1768 (1998).

38. Sussel, L. et al. Mice lacking the homeodomain transcription factor Nkx2.2 have diabetes due to arrested differentiation of pancreatic β cells. Development 125, 2213–2221 (1998).

39. Collombat, P. et al. Opposing actions of Arx and Pax4 in endocrine pancreas development. Genes Dev. 17, 2591–2603 (2003).

40. Dhawan, S., Georgia, S., Tschen, S. I., Fan, G. & Bhushan, A. Pancreatic β Cell Identity Is Maintained by DNA Methylation-Mediated Repression of Arx. Dev. Cell 20, 419–429 (2011).

41. Liu, J. et al. Neurog3-Independent Methylation Is the Earliest Detectable Mark Distinguishing Pancreatic Progenitor Identity. Dev. Cell 48, 49–63 (2019).

42. Papizan, J. B. et al. Nkx2.2 repressor complex regulates islet β-cell specification and prevents β-to-α-cell reprogramming. Genes Dev. 25, 2291–2305 (2011).

43. Wu, X., Li, G. & Xie, R. Decoding the role of TET family dioxygenases in lineage specification. Epigenetics and Chromatin 11, 1–10 (2018).

44. Cirillo, L. A. et al. Opening of compacted chromatin by early developmental transcription factors HNF3 (FoxA) and GATA-4. Mol. Cell 9, 279–89 (2002).

45. Rezania, A. et al. Reversal of diabetes with insulin-producing cells derived in vitro from human pluripotent stem cells. Nat. Biotechnol. 32, 1121–1133 (2014).

46. Zmuda, E. J., Powell, C. A. & Hai, T. A method for murine islet isolation and subcapsular kidney transplantation. J. Vis. Exp. 50, e2096 (2011).

47. Cong, L. et al. Multiplex Genome Engineering Using CRISPR/Cas Systems. Science 339, 819–824 (2013).

48. Wang, X. et al. Noninvasive application of mesenchymal stem cell spheres derived from hESC accelerates wound healing in a CXCL12-CXCR4 axis-dependent manner. Theranostics 9, 6112–6128 (2019).

49. Tu, S. et al. Takusan: A Large Gene Family that Regulates Synaptic Activity. Neuron 55, 69–85 (2007).

50. Dobin, A. et al. STAR: Ultrafast universal RNA-seq aligner. Bioinformatics 29, 15–21 (2013).

51. Anders, S., Pyl, P. T. & Huber, W. HTSeq--a Python framework to work with high-throughput sequencing data. Bioinformatics 31, 166–169 (2015).

52. Love, M. I., Huber, W. & Anders, S. Moderated estimation of fold change and dispersion for RNA-seq data with DESeq2. Genome Biol. 15, 1–21 (2014).

53. Gu, Z., Eils, R. & Schlesner, M. Complex heatmaps reveal patterns and correlations in multidimensional genomic data. Bioinformatics 32, 2847–2849 (2016).

54. Yu, G., Wang, L. G., Han, Y. & He, Q. Y. ClusterProfiler: An R package for comparing biological themes among gene clusters. OMICS 16, 284–287 (2012).

55. Wang, L., Wang, S. & Li, W. RSeQC: Quality control of RNA-seq experiments. Bioinformatics 28, 2184–2185 (2012).

56. Buenrostro, J. D., Wu, B., Chang, H. Y. & Greenleaf, W. J. ATAC-seq: A Method for Assaying Chromatin Accessibility Genome-Wide. Curr. Protoc. Mol. Biol. 109, 21–29 (2015).

57. Quinlan, A. R. & Hall, I. M. BEDTools: A flexible suite of utilities for comparing genomic features. Bioinformatics 26, 841–842 (2010).

58. Villanueva, R. A. M. & Chen, Z. J. ggplot2: Elegant Graphics for Data Analysis. Meas. Interdiscip. Res. Perspect. 17, 160–167 (2019).

59. Heinz, S. et al. Simple Combinations of Lineage-Determining Transcription Factors Prime cis-Regulatory Elements Required for Macrophage and B Cell Identities. Mol. Cell 38, 576–589 (2010).

60. Xi, Y. & Li, W. BSMAP: Whole genome bisulfite sequence MAPping program. BMC Bioinformatics 10, 1–9 (2009).

61. Ramírez, F., Dündar, F., Diehl, S., Grüning, B. A. & Manke, T. DeepTools: A flexible platform for exploring deep-sequencing data. Nucleic Acids Res. 42, 187–191 (2014).

62. Lin, X. et al. BSeQC: Quality control of bisulfite sequencing experiments. Bioinformatics 29, 3227–3229 (2013).

63. Feng, J., Liu, T., Qin, B., Zhang, Y. & Liu, X. S. Identifying ChIP-seq enrichment using MACS. Nat. Protoc. 7, 1728–1740 (2012).

